# Endogenous Rab29 does not impact basal or nigericin and monensin stimulated LRRK2 pathway activity

**DOI:** 10.1101/2020.06.08.139675

**Authors:** Alexia F. Kalogeropulou, Jordana B. Freemantle, Pawel Lis, Edmundo G. Vides, Nicole K. Polinski, Dario R. Alessi

**Affiliations:** Medical Research Council Protein Phosphorylation and Ubiquitylation Unit, School of Life Sciences, University of Dundee, Dow Street, Dundee DD1 5EH, UK; Department of Biochemistry, Stanford University School of Medicine, Stanford, USA 94305-5307; Michael J Fox Foundation for Parkinson’s Research, Grand Central Station, PO Box 4777, New York, NY 10163, U.S.A

## Abstract

Mutations that enhance LRRK2 protein kinase activity cause inherited Parkinson’s disease. LRRK2 phosphorylates a group of Rab GTPase proteins, including Rab10 and Rab12, within the effector-binding switch-II motif. Previous work has indicated that the PARK16 locus, which harbors the gene encoding for Rab29, is involved in Parkinson’s, and that Rab29 operates in a common pathway with LRRK2. Co-expression of Rab29 and LRRK2 stimulates LRRK2 activity by recruiting LRRK2 to the surface of the trans Golgi network. Here we report that knock-out of Rab29 does not influence endogenous LRRK2 activity, based on assessment of Rab10 and Rab12 phosphorylation, in wildtype LRRK2, LRRK2[R1441C] or VPS35[D620N] knock-in mouse tissues and primary cell lines, including brain extracts and embryonic fibroblasts. We find that in brain extracts, Rab12 phosphorylation is more robustly impacted by LRRK2 inhibitors and pathogenic mutations than Rab10 phosphorylation. Transgenic overexpression of Rab29 in a mouse model was also insufficient to stimulate basal LRRK2 activity. We observed that monovalent cation ionophore antibiotics nigericin and monensin enhance LRRK2-mediated Rab10 and Rab12 phosphorylation 4 to 9-fold, in a manner that is independent from Rab29. Moderate stimulation of Rab10 and Rab12 induced by lysosome stressors chloroquine and LLOMe was also not regulated by Rab29. Our findings indicate that basal, pathogenic, as well as nigericin and monensin stimulated LRRK2 pathway activity is not controlled by Rab29. Further work is required to establish how LRRK2 activity is regulated, and whether other Rab proteins can control LRRK2 by targeting it to diverse membranes.

## Introduction

Autosomal dominant missense mutations that hyperactivate LRRK2 (leucine rich repeat kinase 2) are one of the most common causes of familial Parkinson’s disease (PD) [1–4]. Age of onset and progression of LRRK2-driven PD is virtually indistinguishable from sporadic PD, which comprises the vast majority of the patient population. LRRK2 is a large, multi-functional protein kinase that encodes two central catalytic regions, a Roc-type GTPase domain adjacent to a COR (C-terminal of Roc) domain, which is followed by a serine/threonine protein kinase domain. These enzymatic regions are surrounded by several domains, including N-terminal armadillo and ankyrin domains, leucine-rich repeats and a C-terminal WD-40 repeat [5]. The most prevalent LRRK2 pathogenic variants map to either the GTPase Roc [N1437H, R1441C/G/H] or COR [Y1699C] domains, or the kinase domain [G2019S, I2020T], and act as gain-of-function mutations by enhancing LRRK2 kinase activity [6–10]. The mutations within the GTPase Roc/COR domain are proposed to inhibit GTPase activity and enhance GTP binding [11–13]. The mutations within the GTPase domain do not directly activate LRRK2 activity *in vitro*, but enhance interaction with Rab29 located at the Golgi [14,15]. This leads to the recruitment of LRRK2 to the Golgi membrane surface which enhances its kinase activity, as evidenced by increased autophosphorylation at Ser1292 and phosphorylation of its physiological Rab protein substrates, through a yet undefined mechanism [14,16,17]. Mutations within the kinase domain directly stimulate LRRK2 activity by promoting a more closed, active conformation of the catalytic moiety [18–20]. The VPS35[D620N] autosomal dominant mutation that causes PD markedly elevates Rab protein phosphorylation by LRRK2 through an unknown mechanism [21]. LRRK2 is constitutively phosphorylated on several well-studied serine residues in its N-terminus, specifically Ser910, Ser935, Ser955, and Ser973, and these sites are rapidly dephosphorylated upon pharmacological inhibition of LRRK2 [22,23]. LRRK2 protein kinase inhibitors are currently in early stage clinical trials for LRRK2-driven PD [24,25].

Well-characterized and validated substrates of LRRK2 comprise a subset of Rab GTPases that include Rab8A, Rab10, Rab12 and Rab29 [7,26]. Rab GTPases are crucial regulators of intracellular vesicle trafficking, implicated in vesicle formation and transport between target membranes in a tightly controlled network [27]. They influence biology by interacting with specific effector proteins when complexed to GTP. LRRK2 phosphorylates Rab proteins at a highly conserved Ser/Thr residue located at the center of the effector binding region of these enzymes that is also known as the Switch-II motif [7,26,28]. This phosphorylation event appears to act in two ways. Firstly, it prevents Rab proteins interacting with many of their known interactors including guanine nucleotide exchange factors (GEFs) and guanine nucleotide dissociation inhibitors (GDIs) that are required for the shuttling of Rab proteins between membrane compartments. This results in the LRRK2-phosphorylated Rab proteins accumulating on the surface of the compartment on which they are phosphorylated [7,29]. Secondly, LRRK2-phosphorylated Rab8A and Rab10 bind preferentially to a set of effectors, such as RILPL1 and RILPL2, which are implicated in ciliogenesis [26]. These effectors possess a RH2 domain that functions as a phospho-Rab recognition domain [30]. Pathogenic LRRK2 mutants decrease primary cilia formation in cell culture in a manner that is rescued upon LRRK2 inhibition [26]. The ability of LRRK2 to inhibit ciliogenesis requires RILPL1 binding to LRRK2-phosphorylated Rab8A and Rab10 [31,32]. Recent work showed that LRRK2-phosphorylated Rab proteins are dephosphorylated by a highly selective PPM1H protein phosphatase [33].

Significant effort has focused on Rab29, also known as Rab7L1, and its possible roles in regulating LRRK2. The gene encoding Rab29 lies within a genetically complex locus termed PARK16, which is implicated with increased PD risk [34–37]. The PARK16 locus contains 5 genes, and it is not clear which of these genes is relevant for PD or how the numerous variants identified within this locus affect gene expression and/or function. Single nucleotide polymorphisms in non-coding regions of the PARK16 locus have been linked to increasing the transcriptional regulation of Rab29 mRNA [15,38,39]. Several earlier studies alluded to the possibility of Rab29 and LRRK2 acting in converging pathways, by demonstrating epistatic interactions between polymorphisms in the LRRK2 and Rab29 genes that increase PD risk [39,40]. Physical interaction between LRRK2 and Rab29, either *in vitro* or based on a co-immunoprecipitation analysis, has also been demonstrated [15,39,41,42]. Furthermore, analysis of genetic models reveals that Rab29 and LRRK2 operate coordinately to control axon elongation in *C. elegans*, and lysosomal trafficking and kidney pathology in mice [43]. A recent study reports that combined knock-out of LRRK2 and Rab29 does not result in a PD-relevant neuronal pathology or behavioral abnormalities [44]. Rab29 has been implicated in maintaining Golgi morphology and in mediating the retrograde trafficking of the mannose-6-phosphate receptor (M6PR), which recognizes and delivers lysosomal enzymes from the *trans* Golgi to late endosomes and lysosomes [45,46]. An intriguing finding that implicates Rab29 in immune response demonstrated its recruitment to *S. typhi*-containing vacuoles and Rab29 involvement in the generation of typhoid toxin transport intermediates that release the toxin into the extracellular environment [47].

Rab29 belongs to a subfamily of Rab GTPases with Rab32 and Rab38, which are localized to melanosomes and are involved in regulating the trafficking of melanogenic enzymes between the *trans* Golgi and melanosomes [48]. Rab29 is unique among the Rab proteins targeted by LRRK2 in that it possesses two adjacent phosphorylated residues within its Switch-II motif, namely Thr71 and Ser72. Ser72 aligns with the phosphorylation site found in other LRRK2 substrates. To our knowledge there is no evidence that endogenous Rab29 is directly phosphorylated by LRRK2, however in overexpression studies, LRRK2 triggers phosphorylation of both Thr71 and Ser72 in a manner that is blocked with LRRK2 inhibitors [14,26]. Based on mutagenesis overexpression experiments, phosphorylation of these sites on Rab29 was proposed to function as a negative feedback loop to block activation of LRRK2 by Rab29 [14]. Rab32 and Rab38 are not phosphorylated by LRRK2 and do not possess a Ser/Thr residue at the equivalent position within their switch-II motif [26]. Recent work has established that the activation of LRRK2 by Rab29 occurs irrespective of the identity of the membrane to which Rab29 is attached [29], and requires Rab prenylation and nucleotide binding by both LRRK2 and Rab29 [16,29]. Thus, numerous genetic studies in addition to data presented *in vitro*, in cells, and in model organisms, provide substantial evidence that LRRK2 and Rab29 pathways intersect.

Previous work showing that LRRK2 is activated by recruitment to the Golgi *via* interaction with Rab29 is largely based in overexpression experiments in which both Rab29 and wildtype or pathogenic mutants of LRRK2 are co-expressed. In this study, we sought to investigate the physiological relevance of Rab29 as a regulator of endogenous LRRK2 in cell lines as well as in mouse tissues. We describe our efforts to thoroughly characterize four different mouse models that we generated to assess Rab29 impact on basal activity of wildtype and the LRRK2[R1441C] pathogenic mutant. Our data demonstrate that knock-out or moderate transgenic overexpression of Rab29 in a new mouse model we created, does not significantly impact the ability of LRRK2 to phosphorylate Rab10 or Rab12. Furthermore, we show that knock-out of Rab29 does not impact the ability of the VPS35[D620N] mutation or monovalent cation ionophore antibiotics nigericin and monensin to promote Rab10 and Rab12 protein phosphorylation. Our data indicate that Rab29 is not a major regulator of basal wildtype or LRRK2[R1441C] activity as measured in whole cell or tissue extracts that we have analyzed. Further work is therefore required to clarify how LRRK2 activity is regulated.

## Materials and Methods

### Reagents

MLi-2 LRRK2 inhibitor was synthesized by Natalia Shpiro (University of Dundee) and was first described to be a selective LRRK2 inhibitor in previous work [49]. Microcystin-LR was purchased from Enzo Life Sciences (ALX-350-012), oriole fluorescent gel stain was purchased from Bio-Rad (#161-0495), and nigericin was purchased from Invivogen (tlrl-nig). Monensin sodium salt (M5273), Leu-Leu methyl ester hydrobromide (LLOMe) (L7393), and chloroquine diphosphate salt (C6628) were purchased from Sigma Aldrich.

### Generation of MJFF rabbit monoclonal Rab29 total antibodies (MJF-30)

Rabbit immunization and rabbit antibody generation was performed by Abcam Inc. (Burlingame, CA). To generate the Rab29 total antibodies, full length recombinant proteins as well as N-terminal and C-terminal peptides were used. Abcam performed 3 subcutaneous injections using the immunogens conjugated with keyhole limpet haemocyanin (KLH), followed by 2 subcutaneous injections using the immunogens conjugated with ovalbumin. Target immunogen is described in Table 1. Following the initial injections with full-length proteins, booster immunizations using N-terminal and C-terminal peptides were performed. The provided bleeds (1 bleed pre-immunization, 2 bleeds after immunization with full-length protein, 1 bleed after additional immunization with the N- and C-terminal peptides) from immunized animals (6 rabbits – E8767-E8772) were tested at 1:1000 dilution using lysates of A549 wildtype and Rab29 knock-out cell lysates. Rabbits producing the best antibody were chosen for monoclonal antibody generation. Hybridoma fusion was performed according to an established protocol [50]. Abcam used a process omitting the multi-clone stage and provided 81 single clones directly. Single-clone supernatants were screened using immunoblots of A549 wildtype and Rab29 knock-out cell lysates. The top eight clones that gave the most robust signal were additionally tested using MEF lysates to check for human/mouse Rab29 specificity. Clone #124 was able to detect both human and mouse Rab29, while clone #104, which only detected human Rab29, showed the strongest and cleanest signal. Based on these results, clones #104 (human selective catalogue number ab256527) and clone #124 (human + mouse selective antibody catalogue number ab256526) were chosen for recombinant antibody generation and are now commercially available from Abcam.

**Table 1.**
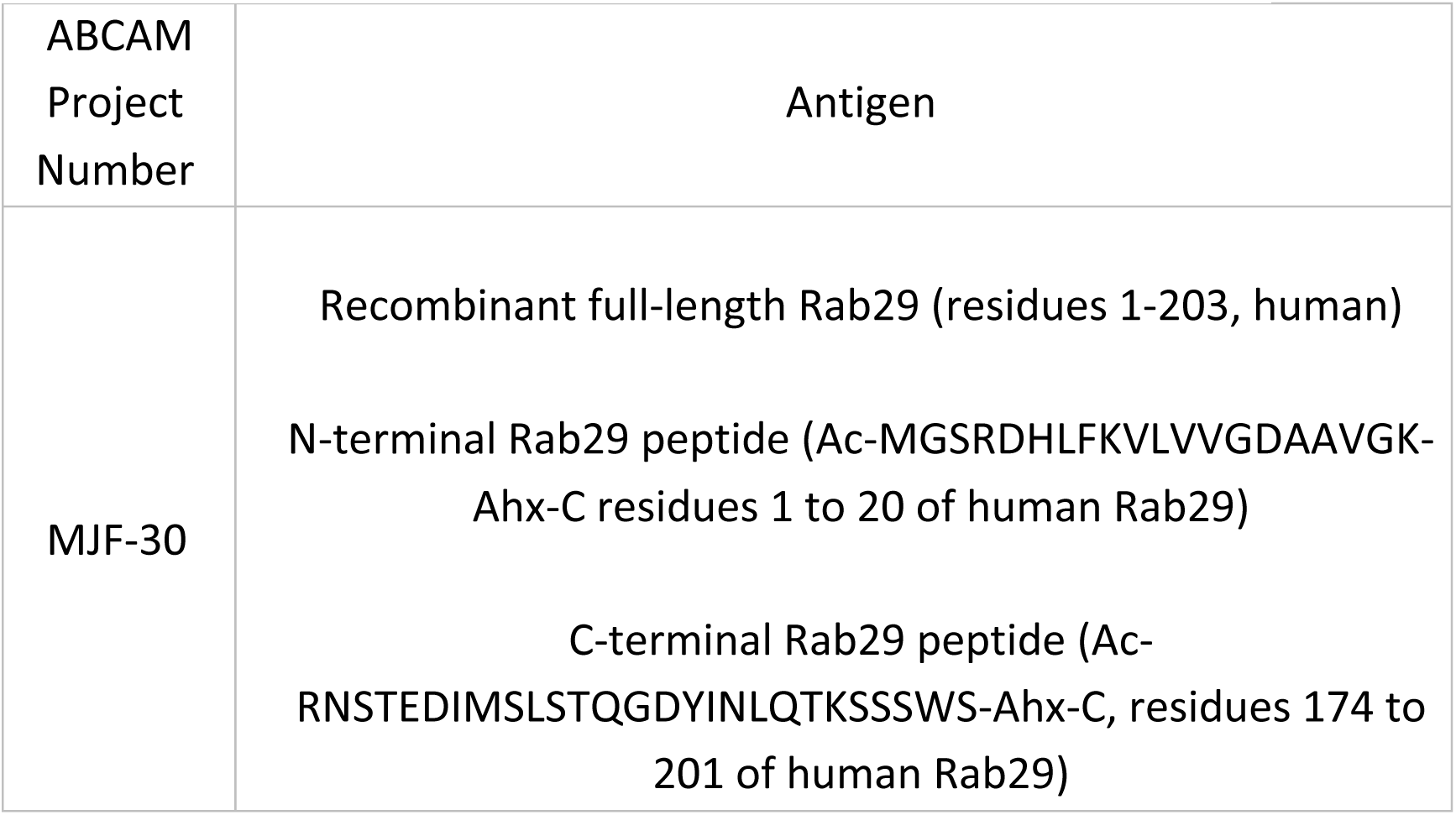
Antigens used in the MJFF rabbit monoclonal Rab29 program.

### Other Antibodies

The MJFF rabbit monoclonal antibody Rab10 pThr73 was previously characterized [51] and purchased through Abcam (ab230261). The MJFF rabbit monoclonal Rab12 pSer105 (equivalent phosphorylation site to human pSer106) antibody was described previously [21] and is available from Abcam (ab256487). Recombinant anti-LRRK2 pSer1292 MJFR-19-7-8 (ab203181), recombinant anti-Rab8A MJF-R22 antibody (ab237702), recombinant anti-M6PR (cation independent) antibody (ab124767) and rabbit polyclonal antibody VDAC1 were also purchased from Abcam (ab15895). The mouse monoclonal antibody against total LRRK2 (C-terminus) was purchased from NeuroMab (clone N241A/34, #75-253). Rabbit monoclonal anti-Rab10 (#8127), mouse monoclonal alpha-tubulin (#3873), and rabbit monoclonal PDI (#3501) were purchased from Cell Signaling Technology. Mouse monoclonal Rab32 (B-4) antibody was purchased from Santa Cruz Biotechnology (sc-390178) and was diluted 1:200. The mouse monoclonal anti-Rab10 total antibody was purchased from Nanotools (#0680–100/Rab10-605B11) and used at a final concentration of 1 μg/ml. Mouse anti-glyceraldehyde-3-phosphate dehydrogenase (GAPDH) was purchased from Santa Cruz Biotechnology (sc-32233) and was used at 1:2000. Mouse monoclonal ACBD3 antibody was purchased from Sigma (WH0064746M1) and used at 10 μg/ml for immunofluorescence. Rabbit monoclonal antibodies for total LRRK2 (N-terminus) (UDD3) and pSer935 LRRK2 (UDD2), and sheep polyclonal antibodies Rab29 total (S984D), Rab12 total (SA227), Rab35 total (SA314) and PPM1H total (DA018) were purified by MRC PPU Reagents and Services at the University of Dundee and were all used at a final concentration of 1 μg/ml. All rabbit and mouse primary antibodies were diluted 1:1000 (unless otherwise stated) in 5% (w/v) bovine serum albumin (BSA) dissolved in TBS-T (50 mM Tris base, 150 mM sodium chloride (NaCl), 0.1% (v/v) Tween 20). Sheep polyclonal antibodies were diluted in 5% (w/v) skim milk powder dissolved in TBS-T. Goat anti-mouse IRDye 800CW (#926-32210), goat anti-mouse IRDye 680LT (#926-68020), goat anti-rabbit IRDye 800CW (#926-32211), and donkey anti-goat IRDye 800CW (#926-32214) IgG (H+L) secondary antibodies were from LI-COR and were diluted 1:10000 in 5% (w/v) milk in TBS-T.

### Quantitative immunoblot analysis

Cell or tissue extracts were mixed with a quarter of a volume of 4x SDS-PAGE loading buffer [250 mM Tris–HCl, pH 6.8, 8% (w/v) SDS, 40% (v/v) glycerol, 0.02% (w/v) bromophenol blue and 5% (v/v) 2-mercaptoethanol] and heated at 95 °C for 5 minutes. Samples ranging from 15-40 μg were loaded onto a NuPAGE 4–12% Bis–Tris Midi Gel (Thermo Fisher Scientific, Cat# WG1402BOX or Cat# WG1403BOX) or self-cast 10% Bis-Tris gel and electrophoresed at 130 V for 2 h with NuPAGE MOPS SDS running buffer (Thermo Fisher Scientific, Cat# NP0001-02). At the end of electrophoresis, proteins were electrophoretically transferred onto a nitrocellulose membrane (GE Healthcare, Amersham Protran Supported 0.45 μm NC) at 90 V for 90 min on ice in transfer buffer (48 mM Tris-HCl and 39 mM glycine supplemented with 20% methanol). The transferred membrane was blocked with 5% (w/v) skim milk powder dissolved in TBS-T (50 mM Tris base, 150 mM sodium chloride (NaCl), 0.1% (v/v) Tween 20) at room temperature for 1 h. Membranes were washed three times with TBS-T and were incubated in primary antibody overnight at 4 °C. Prior to secondary antibody incubation, membranes were washed three times for 15 min with TBS-T. The membranes were incubated with secondary antibody for 1 h at room temperature. Thereafter, membranes were washed with TBS-T three times with a 15 min incubation for each wash, and protein bands were acquired *via* near infrared fluorescent detection using the Odyssey CLx imaging system and quantified using the Image Studio software.

### Mice

Mice selected for this study were maintained under specific pathogen-free conditions at the University of Dundee (U.K.). All animal studies were ethically reviewed and carried out in accordance with the Animals (Scientific Procedures) Act 1986 and regulations set by the University of Dundee and the U.K. Home Office. Animal studies and breeding were approved by the University of Dundee ethical committee and performed under a U.K. Home Office project license. Mice were housed at an ambient temperature (20–24 °C) and humidity (45– 55%) and were maintained on a 12 h light/12 h dark cycle, with free access to food (SDS RM No. 3 autoclavable) and water. For the experiments described in Fig 2B-D and Fig S1 (Rab29 knock-out model), Fig 5A-F (transgenic Rab29 overexpression model), Fig 6B-E and Fig S4 (Rab29 knock-out LRRK2[R1441C] knock-in model), and Fig 7B-E and Fig S6 (Rab29 knock-out VPS35[D620N] knock-in model), 6-month-old littermate matched or matched mice of the indicated genotypes were injected subcutaneously with vehicle [40% (w/v) (2-hydroxypropyl)-β-cyclodextrin (Sigma–Aldrich #332607)] or MLi-2 dissolved in vehicle at a 30 mg/kg final dose. Mice were killed by cervical dislocation 2 h following treatment and the collected tissues were rapidly snap-frozen in liquid nitrogen. For the experiment outlined in Fig S2 (transgenic Rab29 overexpression model), mice ranging from 3 to 4 months and of the indicated genotypes were dosed, killed and the tissues were isolated as outlined above. For the studies involving 18-24 mice which took place over multiple days, the genotypes and treatments were randomized each day to account for any temporal differences, such as light and ventilation. A general overview of the mouse models utilized in this study is outlined in Table 2.

**Table 2.** Summary of mouse models used in this study.

| Name of model | Strain designation | Provider | Previously characterized |
| --- | --- | --- | --- |
| Rab29 knock-out | Rab29<sup>tm1a(EUCOMM)Wtsi</sup>/WtsiOulu | Wellcome Sanger Institute, Infrafrontier EMMA mouse repository<br>EM: 05517 | [43,44,46] |
| LRRK2[R1441C] | B6.Cg-Lrrk2<sup>tm1.1Shn</sup>/J | Jackson Laboratories<br>Stock No: 009346 | [26,51] |
| VPS35[D620N] | B6(Cg)-Vps35<sup>tm1.1Mjff</sup>/J | Jackson Laboratories<br>Stock No: 023409 | [21] |
| Transgenic Rab29 overexpression | C57BL/6NTac-Gt(ROSA)26Sor<sup>tm1(Pgk-Rab29)Tac</sup> | Taconic<br>Model No: 16552 |  |

**Figure 1.**
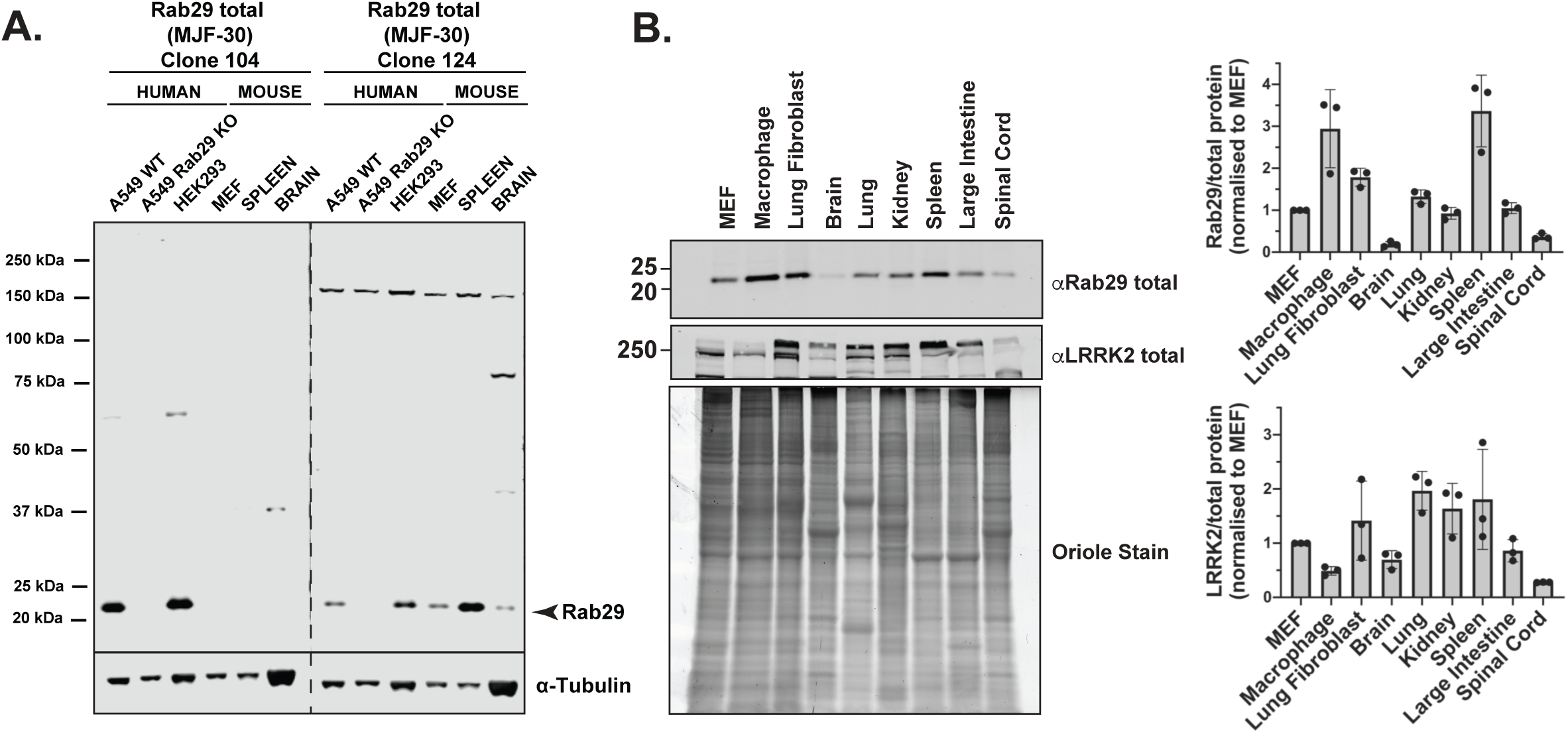
Detection of endogenous Rab29 in human and mouse extracts with novel Rab29 monoclonal antibodies. **(A)** 30 μg of the indicated cell and tissue extracts were subjected to quantitative immunoblot analysis with the indicated antibodies, diluted to 1 μg/ml. Membranes were developed using the LI-COR Odyssey CLx Western Blot imaging system. **(B)** Various cell and tissue extracts were analyzed for Rab29 and LRRK2 expression (upper panels). Total protein levels were visualized using the Bio-Rad Oriole Fluorescent Gel Stain and exposed using the Bio-Rad Chemidoc MP Imaging System (lower panel). Total protein levels were quantified using Bio-Rad Image Lab software. Total Rab29 or total LRRK2 was quantified using the Image Studio software. Quantified data are presented as ratios of Rab29 or LRRK2 expression divided by total protein levels, and values were normalized to the average of the respective protein expression observed in MEFs. Quantifications are representative of three independent experiments and shown as mean ± SD.

**Figure 2.**
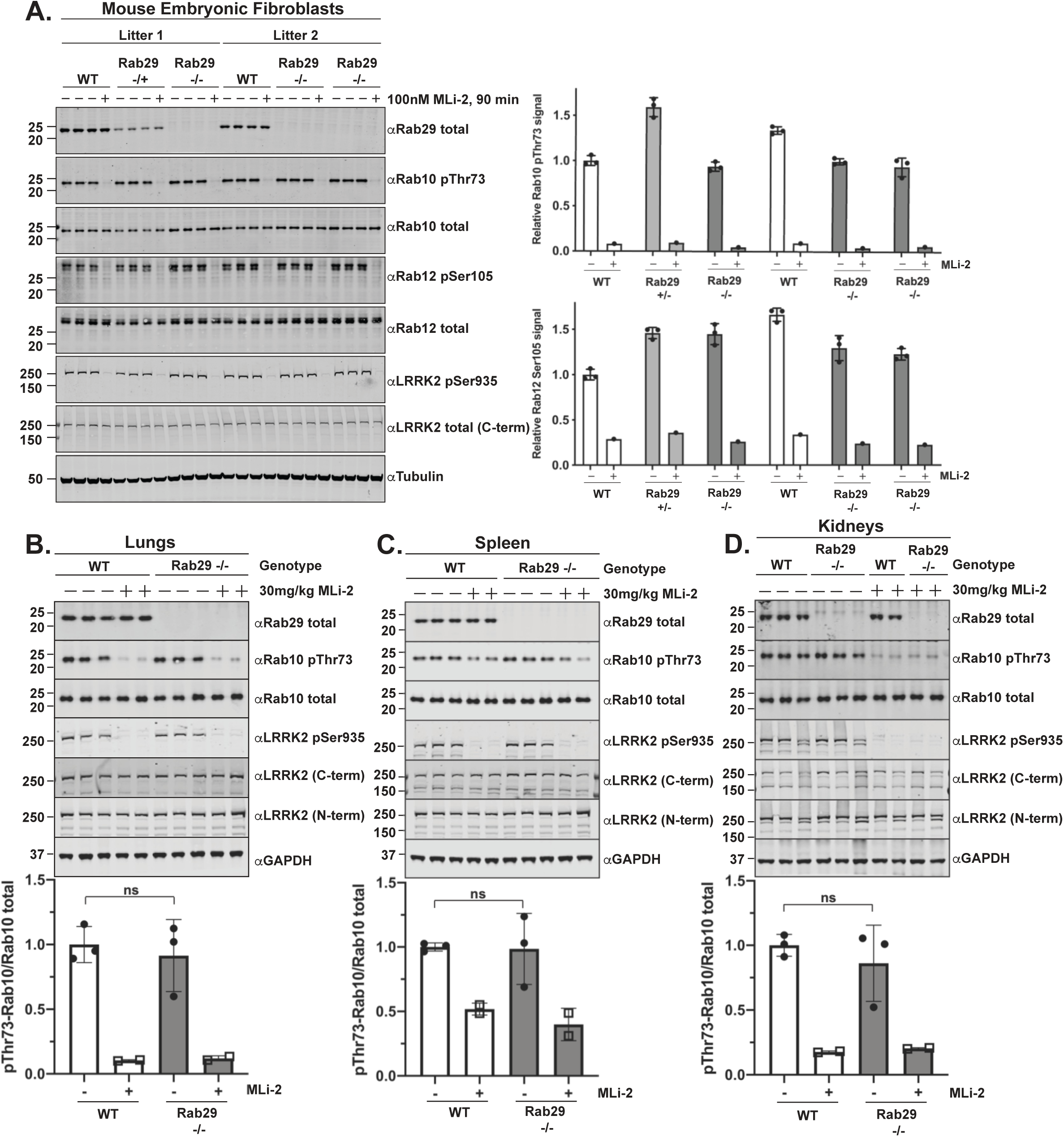
Knock-out of Rab29 does not affect basal LRRK2 activity. **(A)** Two independent littermate-matched wildtype (WT), Rab29 knock-out heterozygous (+/-), or Rab29 knock-out homozygous (-/-) MEFs were treated with vehicle (DMSO) or 100 nM LRRK2 inhibitor MLi-2 for 90 min prior to harvest. 20 μg of whole cell extracts were subjected to quantitative immunoblot analysis with the indicated antibodies. Technical replicates represent cell extract obtained from a different dish of cells. The membranes were developed using the LI-COR Odyssey CLx Western Blot imaging system. Quantified data are presented as the mean ± SD of phospho-Rab10/total Rab10 and phospho-Rab12/total Rab12 ratios, which were quantified using the Image Studio software. Values were normalized to the average of Litter 1 wildtype MEFs treated with DMSO. Similar results were obtained in three independent experiments. **(B-D)** 6-month-old, littermate-matched wildtype (WT) and Rab29 knock-out (-/-) mice were administered with vehicle (40% (w/v) (2-hydroxypropyl)-β-cyclodextrin) or 30 mg/kg MLi-2 dissolved in vehicle by subcutaneous injection 2 h prior to tissue collection. 40 μg of tissue extracts were analyzed by quantitative immunoblot as described in (A). Each lane represents tissue extract derived from a different mouse. Quantified data are presented as mean ± SD and values were normalized to the average of the wildtype, vehicle treated mice. Data were analyzed using two-tailed unpaired t-test and there was no statistical significance between WT and Rab29 knock-out mice. The resulting P-values from the unpaired t-tests are (B) lungs P = 0.6585 (C) spleen P = 0.9318 (D) kidneys P = 0.4793.

**Figure 3.**
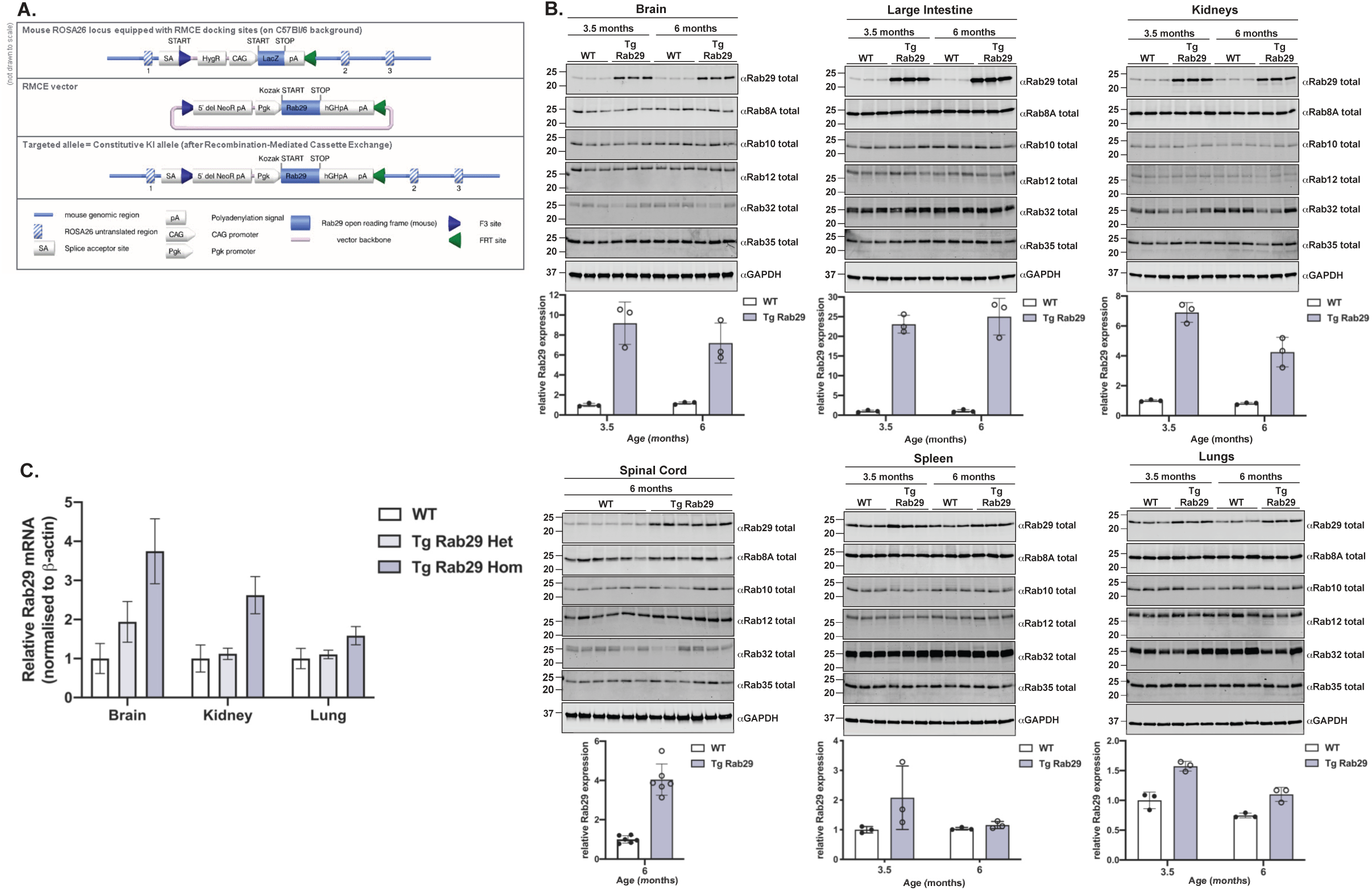
Characterization of MJFF transgenic Rab29 overexpressing mouse. **(A)** Schematic overview of design and targeting strategy utilized to create the Rab29 transgenic mouse model. The illustration depicts the transgene containing a neo cassette, a Pgk promoter, the murine Rab29 open reading frame together with a Kozak sequence (GCCACC), the human growth hormone polyadenylation signal and an additional polyadenylation signal. These elements were inserted into the ROSA26 locus *via* recombinase-mediated cassette exchange. **(B)** The indicated tissues were collected from 3.5-month-old and 6-month-old wildtype (WT) and homozygous, transgenic Rab29 overexpressing (Tg Rab29) mice. 30 μg of tissue extracts were subjected to quantitative immunoblot analysis with the indicated antibodies. The membranes were developed using the LI-COR Odyssey CLx Western Blot imaging system. Quantified data are presented as mean ± SD of total Rab29/GAPDH ratios, calculated using the Image Studio software. Values were normalized to the average of the 3.5-month-old wildtype mice. Each lane represents a tissue sample from a different animal. **(C)** The indicated tissue extracts from 3.5-month-old wildtype (WT), heterozygous (Tg Rab29 Het), and homozygous (Tg Rab29 Hom) transgenic Rab29 overexpressing mice were processed for total RNA extraction, cDNA synthesis, and subsequent quantitative real time RT-PCR analysis. Rab29 mRNA levels were normalized to β-actin mRNA levels and further normalized to the wildtype of the appropriate tissue. Quantified data are presented as mean ± SD of four biological replicates per genotype.

**Figure 4.**
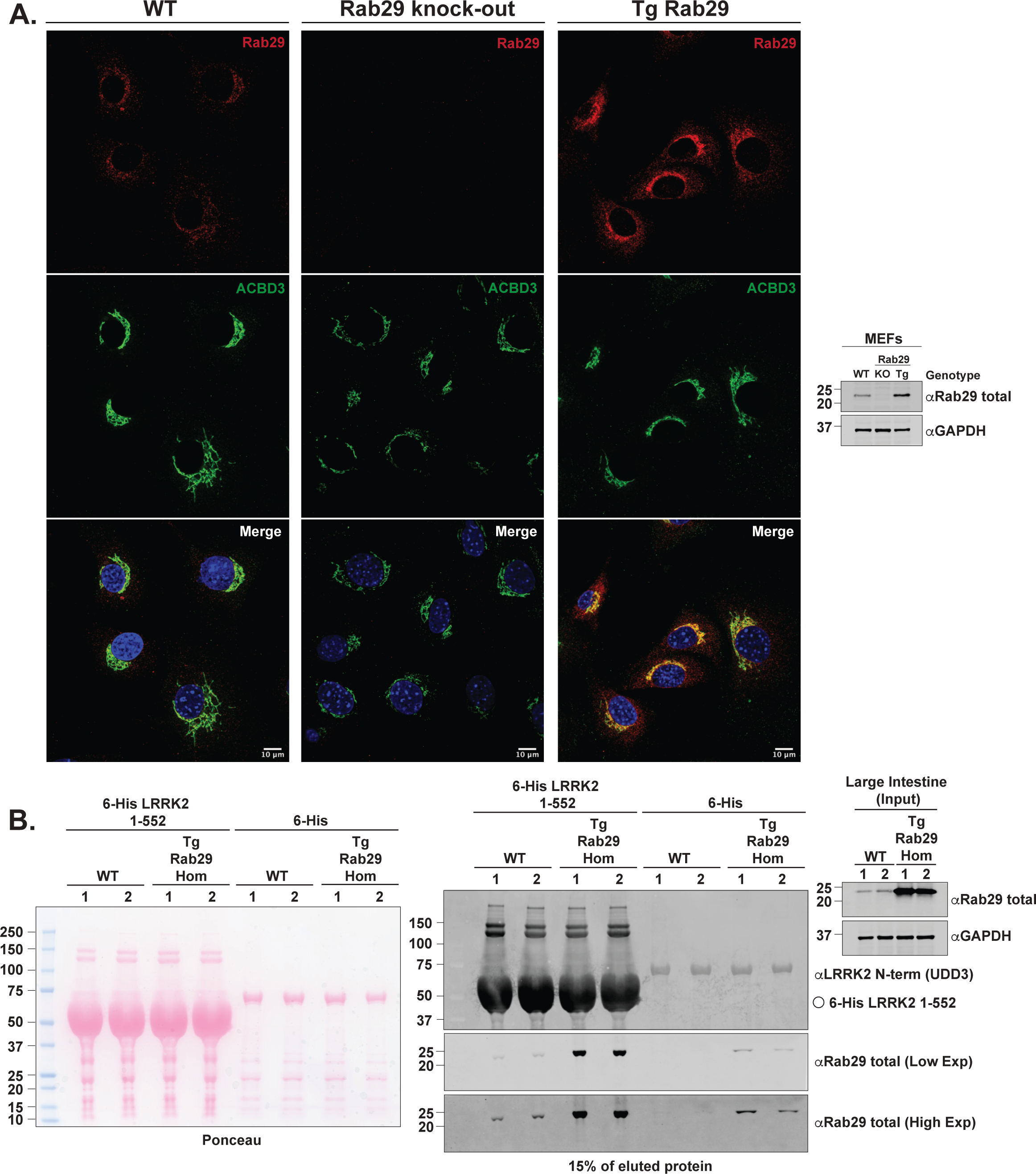
Transgenic Rab29 in overexpressing mouse is functional and competent for binding GTP. **(A)** Littermate matched wildtype and transgenic Rab29 overexpressing MEFs, and Rab29 knock-out MEFs were seeded on coverslips. The following day, cells were fixed with 4% (v/v) paraformaldehyde and Rab29 was visualized with rabbit Rab29 antibody and anti-rabbit Alexa Fluor 594 secondary. Golgi were stained with mouse anti-ACBD3 and anti-mouse Alexa Fluor 488 secondary, and nuclei using DAPI. Immunoblotting of MEFs of the indicated genotypes was undertaken in parallel. **(B)** Recombinant 6-His LRRK2 1-552 immobilized on nickel-NTA resin or immobilized 6-His, was incubated with intestinal tissue extracts supplemented with excess GTP-γ-S from two different wildtype or transgenic Rab29 overexpressing mice. Eluted protein was analyzed by immunoblotting with the indicated antibodies. The membranes were developed using the LI-COR Odyssey CLx Western Blot imaging system. Similar results were obtained in two independent experiments.

**Figure 5.**
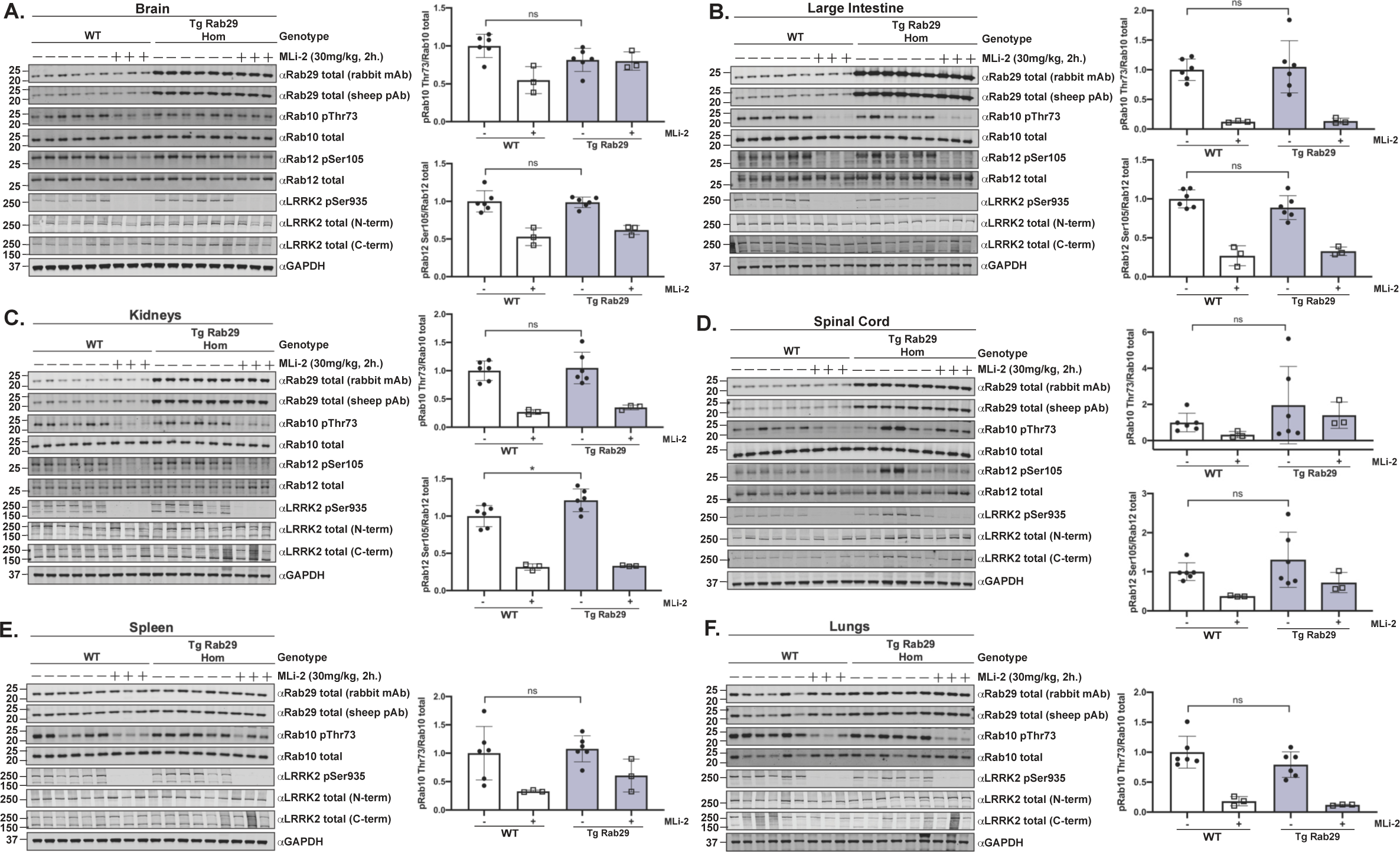
Overexpression of Rab29 does not impact LRRK2-mediated Rab10 phosphorylation. **(A-F)** 6-month-old littermate-matched wildtype (WT) and homozygous, transgenic Rab29 overexpressing (Tg Rab29 Hom) mice were administered with vehicle (40% (w/v) (2-hydroxypropyl)-β-cyclodextrin) or 30 mg/kg MLi-2 dissolved in vehicle by subcutaneous injection 2 h prior to tissue collection. 40 μg of tissue extracts were subjected to quantitative immunoblot analysis with the indicated antibodies. The membranes were developed using the LI-COR Odyssey CLx Western Blot imaging system. Quantified data are presented as the ratios of phospho-Rab10/total Rab10 and phospho-Rab12/total Rab12, calculated using the Image Studio software. Values were normalized to the average of the wildtype, vehicle treated mice. Each lane represents a tissue sample from a different animal. Quantifications are presented as mean ± SD and data were analyzed using two-tailed unpaired t-test. There was no statistical significance in Rab10 phosphorylation between wildtype and homozygous, transgenic Rab29 overexpressing mice. The resulting P-values from the aforementioned statistical analyses of Rab10 phosphorylation between genotypes are as follows: (A) brain P = 0.0625 (B) large intestine P = 0.8026 (C) kidneys P = 0.7243 (D) spinal cord P = 0.3120 (E) spleen P = 0.7257 (F) lungs P = 0.1676. The resulting P-values from the statistical analyses of Rab12 phosphorylation between genotypes are: (A) brain P = 0.8545 (B) large intestine P = 0.1831 (C) kidneys *P = 0.0316 (D) spinal cord P = 0.3361.

**Figure 6.**
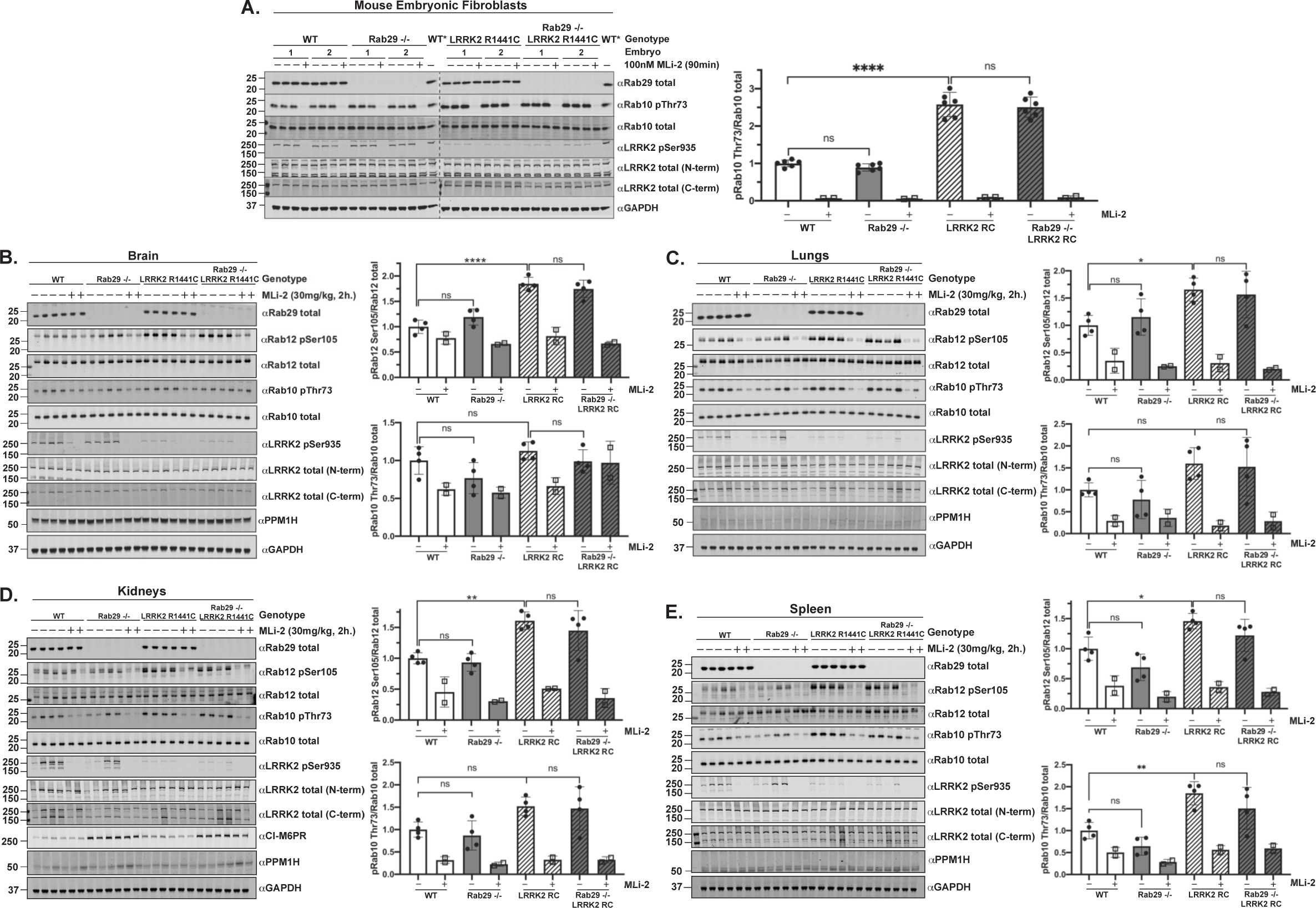
Knock-out of Rab29 does not reduce elevated Rab10 phosphorylation in pathogenic LRRK2[R1441C] knock-in mice. **(A)** The indicated matched primary MEFs were treated with vehicle (DMSO) or 100 nM LRRK2 inhibitor MLi-2 for 90 min prior to harvest. 20 μg of whole cell extracts were subjected to quantitative immunoblot analysis with the indicated antibodies. Technical replicates represent cell extract obtained from a different dish of cells. Cell extract derived from a wildtype MEF cell line (WT*) was added to each gel in order to accurately compare the Rab10 pThr73/Rab10 total ratios between genotypes. The membranes were developed using the LI-COR Odyssey CLx Western Blot imaging system. Quantified data are presented as the ratios of phospho-Rab10/total Rab10, calculated using the Image Studio software. Values were normalized to the average of wildtype MEFs treated with DMSO. Similar results were obtained in two separate experiments. Quantifications are presented as mean ± SD. Data were analyzed by one-way ANOVA with Tukey’s multiple comparisons test and there was a statistically significant difference between wildtype and LRRK2[R1441C] knock-in MEFs (****P < 0.0001) but not between wildtype and Rab29 knock-out MEFs (P = 0.8351) or between LRRK2[R1441C] knock-in and Rab29 knock-out LRRK2[R1441C] knock-in MEFs (P = 0.9452). **(B-D)** The indicated 6-month-old matched mice were administered with vehicle (40% (w/v) (2-hydroxypropyl)-β-cyclodextrin) or 30 mg/kg MLi-2 dissolved in vehicle by subcutaneous injection 2 h prior to tissue collection. 40 μg of tissue extracts were subjected to quantitative immunoblot analysis with the indicated antibodies. The membranes were developed using the LI-COR Odyssey CLx Western Blot imaging system. Quantified data are presented as the ratios of phospho-Rab10/total Rab10 and phospho-Rab12/total Rab12, which were calculated using the Image Studio software. Quantifications are presented as mean ± SD, normalized to vehicle treated, wildtype animals. Each lane represents a tissue sample from a different animal. Data was analyzed by one-way ANOVA with Tukey’s multiple comparisons test and there was a statistically significant difference in Rab10 phosphorylation between wildtype and LRRK2[R1441C] spleen samples (**P = 0.0093 (E)). All other phospho-Rab10 comparisons were not statistically significant. Wildtype vs LRRK2[R1441C]: P = 0.7169 (B), P = 0.2792 (C), P = 0.1555 (D). Wildtype vs Rab29 knock-out: P = 0.2594 (B), P = 0.8942 (C), P = 0.9372 (D), P = 0.3964 (E). LRRK2[R1441C] vs Rab29 knock-out LRRK2[R1441C]: P = 0.6629 (B), P = 0.9955 (C), P = 0.9962 (D), P = 0.4184 (E). There was a statistically significant difference in Rab12 phosphorylation between wildtype and LRRK2[R1441C] tissue samples: brain ****P < 0.0001 (B), lungs *P = 0.0438 (C), kidneys **P = 0.0041 (D), and spleen *P = 0.0390 (E). All other phospho-Rab12 comparisons were not statistically significant. Wildtype vs Rab29 knock-out: P = 0.3057 (B), P = 0.8915 (C), P = 0.9595 (D), P = 0.2078 (E). LRRK2[R1441C] vs Rab29 knock-out LRRK2[R1441C]: P = 0.7791 (B), P = 0.9747 (C), 0.6681 (D), P = 0.4093 (E).

**Figure 7.**
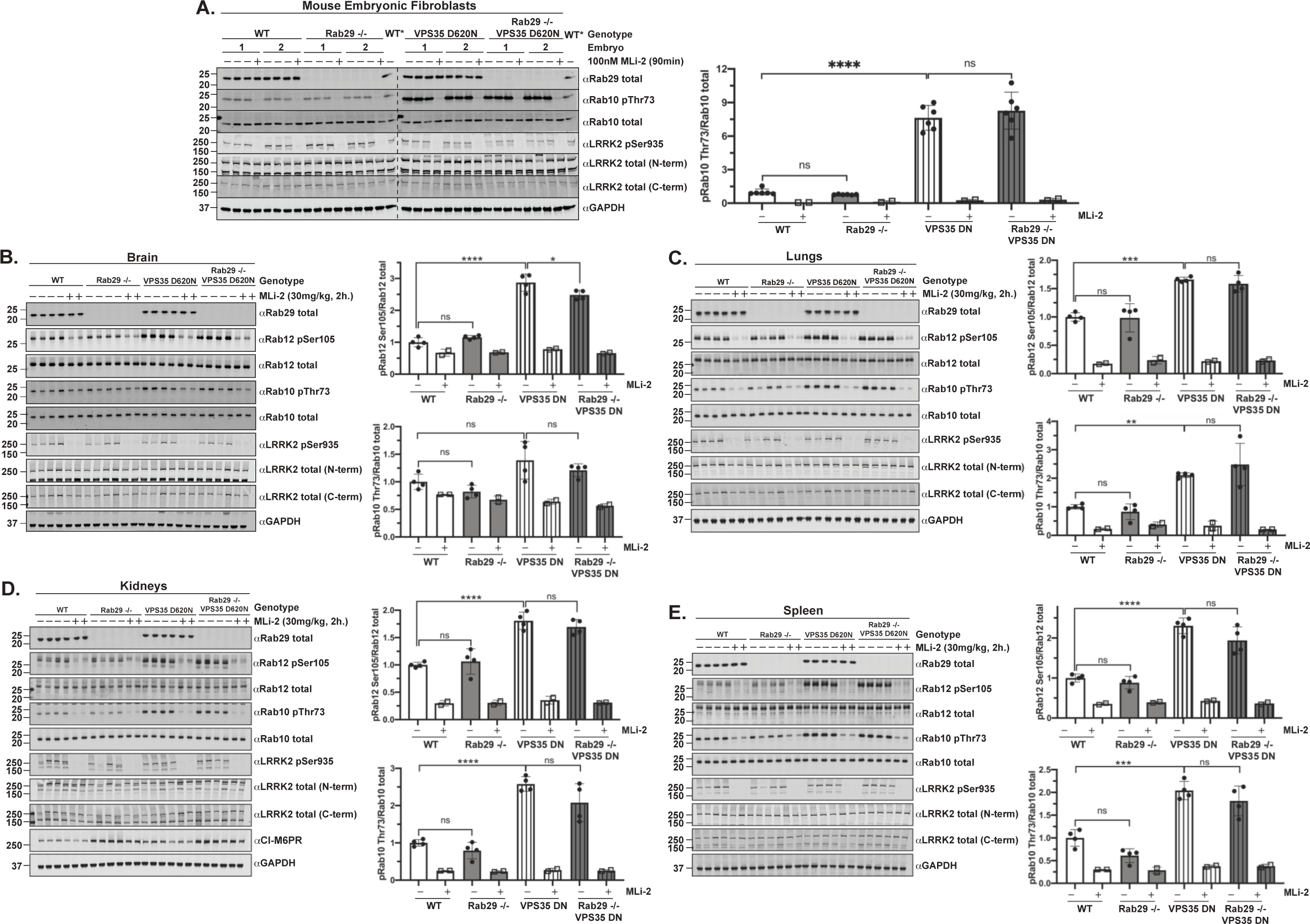
Rab29 knock-out does not reduce the enhanced LRRK2-mediated phosphorylation of Rab10 in VPS35[D620N] knock-in mice. **(A)** The indicated matched MEFs were treated with vehicle (DMSO) or 100 nM LRRK2 inhibitor MLi-2 for 90 min prior to harvest. 15 μg of whole cell extracts were subjected to quantitative immunoblot analysis with the indicated antibodies. Technical replicates represent cell extract obtained from a different dish of cells. Cell extract derived from a wildtype MEF cell line (WT*) was added to each gel in order to accurately compare the Rab10 pThr73/Rab10 total ratios between genotypes. The membranes were developed using the LI-COR Odyssey CLx Western Blot imaging system. Quantified data are presented as the ratios of phospho-Rab10/total Rab10 and phospho-Rab12/total Rab12, calculated using the Image Studio software. Values were normalized to the average of wildtype MEFs treated with DMSO. Similar results were obtained in two separate experiments. Quantifications are presented as mean ± SD. Data were analyzed by one-way ANOVA with Tukey’s multiple comparisons test and there was a statistically significant difference between wildtype and VPS35[D620N] knock-in MEFs (****P < 0.0001) but not between wildtype and Rab29 knock-out MEFs (P = 0.9792) or between VPS35[D620N] knock-in and Rab29 knock-out VPS35[D620N] knock-in MEFs (P = 0.6952). **(B-D)** The indicated 6-month-old matched mice were administered with vehicle (40% (w/v) (2-hydroxypropyl)-β-cyclodextrin) or 30 mg/kg MLi-2 dissolved in vehicle by subcutaneous injection 2 h prior to tissue collection. 40 μg of tissue extracts were subjected to quantitative immunoblot analysis with the indicated antibodies. The membranes were developed using the LI-COR Odyssey CLx Western Blot imaging system. Quantified data are presented as the ratios of phospho-Rab10/total Rab10 and phospho-Rab12/total Rab12, calculated using the Image Studio software. Quantifications are presented as mean ± SD, normalized to vehicle treated, wildtype animals. Each lane represents a tissue sample from a different animal. Data were analyzed by one-way ANOVA with Tukey’s multiple comparisons test and there was a statistically significant difference in Rab10 phosphorylation between wildtype and VPS35[D620N] lungs **P = 0.0094 (C), kidneys ****P < 0.0001 (D), spleen ***P = 0.0001 (E), but not brain (P = 0.0738 (B)). All other phospho-Rab10 comparisons were not statistically significant. Wildtype vs Rab29 knock-out: P = 0.6120 (B), P = 0.9270 (C), P = 0.7648 (D), P = 0.1233 (E). VPS35[D620N] vs Rab29 knock-out VPS35[D620N]: P = 0.5654 (B), P = 0.1329 (C), P = 0.5075 (D). There was a statistically significant difference in Rab12 phosphorylation between wildtype and VPS35[D620N] tissue samples: brain **** P < 0.0001 (B), lungs *** P = 0.0002 (C), kidneys **** P < 0.0001 (D), and spleen **** P < 0.0001 (E). There was a statistically significant difference in Rab12 phosphorylation between VPS35[D620N] and Rab29 knock-out VPS35[D620N] brain samples, *P = 0.0346. All other phospho-Rab12 comparisons were not statistically significant. Wildtype vs Rab29 knock-out: P = 0.5995 (B), P = 0.9988 (C), P = 0.9400 (D), P = 0.8677 (E). VPS35[D620N] vs Rab29 knock-out VPS35[D620N]: P = 0.6015 (B), P = 0.8935 (C), P = 0.7533 (D), P = 0.1332 (E).

### Generation of Rab29 knock-out mice

Rab29 knock-out mice are made available through the Wellcome Trust Sanger Institute, distributed by Infrafrontier: EMMA mouse repository (EM: 05517), and were characterized previously [43]. LacZ-knock-in Rab29^tm1a(EUCOMM)Wtsi^ mice were bred with Taconic Total Body Cre mice expressing Cre recombinase (Model 12524), which recognizes the loxP sites that flank the inserted promoter-driven neomycin cassette and exon 4 of Rab29, which is critical for expression. Following the deletion of exon 4, the mice were then bred and maintained on a C57Bl/6j background to remove the Cre recombinase allele, and to produce the experimental animals used in this study. The genotypes of the Rab29 knock-out mice were confirmed by PCR using genomic DNA isolated from ear biopsies and primers that amplify the entire Rab29 gene, as well as by immunoblotting.

### Generation of Rab29 knock-out LRRK2[R1441C] and Rab29 knock-out VPS35[D620N] knock-in mice

The generation of LRRK2[R1441C] knock-in mice and VPS35[D620N] knock-in mice were described previously [21,26]. Rab29 knock-out heterozygous mice were crossed with LRRK2[R1441C] or VPS35[D620N] knock-in homozygous mice to produce double heterozygous mice for Rab29 knock-out and LRRK2[R1441C] or VPS35[D620N] knock-in. Double heterozygous mice were crossed to produce the following genotypes: Rab29 wildtype/LRRK2 (or VPS35) wildtype, Rab29 knock-out/LRRK2 (or VPS35) wildtype, Rab29 wildtype/LRRK2[R1441C] (or VPS35[D620N]), or Rab29 knock-out/LRRK2[R1441C] (or VPS35[D620N]) (1/16 frequency of indicated genotypes by Mendelian inheritance). Double homozygous mice of the aforementioned genotypes were then expanded as four different subsets to produce additional double homozygous mice for mouse embryonic fibroblast generation and MLi-2 injection studies. Matched mice of the same generation were used for experimental studies.

### Generation of transgenic Rab29 overexpressing mice

The Michael J. Fox Foundation for Parkinson’s Research generated the transgenic Rab29-overexpressing mouse model (C57BL/6NTac-Gt(ROSA)26Sortm1(Pgk-Rab29)Tac), which is made available through Taconic (Model 16552). The constitutive knock-in of Pgk-Rab29 in the ROSA26 locus *via* targeted transgenesis was undertaken by Taconic. The Rab29 sequence was synthesized according to the NCBI transcript NM_144875.2. The following elements were inserted into the ROSA26 locus (NCBI gene ID: 14910) using Recombination-Mediated Cassette Exchange (RMCE): a Pgk promoter, the Rab29 open reading frame together with a Kozak sequence (GCCACC), the human Growth Hormone (hGH) polyadenylation signal and an additional polyadenylation signal. The RMCE vector was transfected into the Taconic Biosciences C57Bl/6 embryonic stem (ES) cell line equipped with RMCE docking sites in the ROSA26 locus. The ES cell line pre-equipped with F3/FRT - RMCE docking sites was grown on a mitotically inactivated feeder layer comprised of mouse embryonic fibroblasts in ES cell culture medium containing Leukemia inhibitory factor and Fetal Bovine Serum. The cells were co-transfected with the circular exchange vector containing the transgene and the recombinase pCAG-Flpe pA. The transfection was performed *via* lipofection with a commercially available kit. From day 2 onwards, the medium was replaced daily with medium containing the appropriate selection antibiotics. The recombinant clones were isolated using positive neomycin resistance selection. On day 7 after transfection, resistant ES cell colonies (ES clones) with a distinct morphology were isolated. The clones were expanded and frozen in liquid nitrogen after extensive molecular validation by Southern Blotting and/or PCR. Quality control of the ES cell line used for transfection was performed at Chrombios GmbH (Germany). Karyotype analysis was undertaken by multicolor fluorescence in situ hybridization with probes for all murine chromosomes (mFISH) to confirm that the cell line meets the quality standard for parental ES cell lines used for targeting experiments. In order to generate chimeras, superovulated BALB/c females were mated with BALB/c males following hormone administration. Blastocysts were isolated from the uterus at dpc 3.5. For microinjection, blastocysts were placed in a drop of DMEM with 15% FCS under mineral oil. A flat tip, piezo actuated microinjection-pipette with an internal diameter of 12-15 micrometer was used to inject 10-15 targeted C57BL/6NTac ES cells into each blastocyst. After recovery, 8 injected blastocysts were transferred to each uterine horn of 2.5 days post coitum, pseudopregnant NMRI females. Chimerism was measured in chimeras (G0) by coat color contribution of ES cells to the BALB/c host (black/white). Germline transmission occurred during an IVF expansion using chimeric males and C57BL/6NTac oocyte donors. The colony is maintained by mating wildtype C57BL/6NTac females to heterozygous males and heterozygous females to wildtype C57BL/6NTac males.

### Mouse genotyping

Genotyping of mice was performed by the MRC genotyping team at the MRC-PPU, University of Dundee, by PCR using genomic DNA isolated from ear biopsies. For this purpose, Primer 1 (5’ CACACACATGGTACACAGATATACATGTAGG 3’) Primer 2 (5’ ACATCCATGACACGACTCTACTATAGAGAT 3’) and Primer 3 (5’ CTATCCCGACCGCCTTACTGC 3’) were used to distinguish between wildtype and Rab29 knock-out alleles (63 °C annealing temp). The VPS35[D620N] knock-in mouse strain required Primer 1 (5’ TCATTCTGTGGTTAGTTCAGTTGAG 3’), Primer 2 (5’ CCTCTAACAACCAAGAGGAACC 3’), and Primer 3 (5’ ATTGCATCGCATTGTCTGAG 3’) to distinguish wildtype from D620N knock-in alleles (60 °C annealing temp). The LRRK2[R1441C] knock-in mouse strain required Primer 1 (5’ CTGCAGGCTACTAGATGGTCAAGGT 3’) and Primer 2 (5’ CTAGATAGGACCGAGTGTCGCAGAG 3’) to identify wildtype and R1441C knock-in alleles (60 °C annealing temp). To detect the constitutive KI allele in the transgenic Rab29 overexpression mouse, Primer 1 (5’ TTGGGTCCACTCAGTAGATGC 3’) and Primer 2 (5’ CATGTCTTTAATCTACCTCGATGG 3’) as well as internal PCR control Primer 1 (5’ GTGGCACGGAACTTCTAGTC 3’) and Primer 2 (5’ CTTGTCAAGTAGCAGGAAGA 3’) were used (58 °C annealing temp). To detect the wildtype allele in the transgenic Rab29 overexpression model, Primer 1 (5’ CTCTTCCCTCGTGATCTGCAACTCC 3’) and Primer 2 (5’ CATGTCTTTAATCTACCTCGATGG 3’) and internal PCR control Primer 1 (5’ GAGACTCTGGCTACTCATCC 3’) and Primer 2 (5’ CCTTCAGCAAGAGCTGGGGAC 3’) were used (58 °C annealing temp). All genotyping primers were used at a final concentration of 10 pmol/μl. PCR reactions were set up and run using KOD Hot Start Polymerase standard protocol. PCR bands were visualized on Qiaexcel (Qiagen) using the standard DNA screening kit cartridge.

### Preparation of mouse tissue lysates

Mouse tissues were collected and snap frozen in liquid nitrogen. Snap frozen tissues were weighed and quickly thawed on ice in a 10-fold volume excess of ice-cold lysis buffer containing 50 mM Tris/HCl pH 7.4, 1 mM EGTA, 10 mM 2-glycerophosphate, 50 mM sodium fluoride, 5 mM sodium pyrophosphate, 270 mM sucrose, supplemented with 1 μg/ml microcystin-LR, 1 mM sodium orthovanadate, complete EDTA-free protease inhibitor cocktail (Roche), and 1% (v/v) Triton X-100. Tissue was homogenized using a POLYTRON homogenizer (KINEMATICA), employing 3 rounds of 10 s homogenization with 10 s intervals on ice. Lysates were centrifuged at 20800 g for 30 min at 4 °C and supernatant was collected for subsequent Bradford assay and immunoblot analysis.

### Quantitative real time RT-PCR analysis of Rab29 mRNA from mouse tissue

Frozen tissue samples were ground to a fine powder in a vessel submerged in liquid nitrogen using a mallet. Powdered tissue (∼5 mg) was dissolved in lysis buffer provided by RNeasy micro kit (Qiagen), supplemented with 1% (v/v) β-mercaptoethanol. Tissue was homogenized in lysis buffer using IKA VIBRAX VXR basic orbital shaker for 5 min at 4 °C at 1000 rpm and total RNA was extracted from tissue lysate following RNeasy micro kit instructions. cDNA was synthesized from total RNA extracts using Bio-Rad iScript cDNA synthesis kit (#170-8891), using a starting template of 150 ng total RNA. The Bio-Rad Sso EvaGreen Supermix (#1725201) was used to set up qPCR reactions in a 384-well plate format according to the manufacturer’s instructions and 20 μl reactions were prepared in duplicate. Real-time quantitative PCR primers for mouse Rab29 were designed using NCBI Primer Blast and the sequences are as follows: 5’-AGGCCATGAGAGTCCTCGTT-3’ (forward) and 5’-GGGCTTGGCTTGGAGATTTGA-3’ (reverse). The β-actin internal control primers employed for real-time quantitative PCR analysis are described in [52]: 5′-CACTATCGGCAATGAGCGGTTCC-3′ (forward) and 5′-CAGCACTGTGTTGGCATAGAGGTC-3′ (reverse). Primers were ordered from Sigma as lyophilized and reconstituted in Milli-Q water to a stock concentration of 100 μM and further diluted to a 10 μM working stock. The PCR efficiency of the primers was validated using the relative standard curve method. The qPCR reactions were run using the Bio-Rad CFX384 Real-Time System C1000 Thermal Cycler, and raw data was collected for duplicate reactions from four biological replicates per genotype for three different tissues. The relative quantification of Rab29 and β-actin mRNA was undertaken using the comparative Ct (cycle threshold) method, which employs the formula RQ = 2^(-ΔΔCt)^ [53].

### LRRK2 fragment purification and Rab29 pulldown assay

pQE80L 2X 6-His LRRK2 1-552 and pET15D 6-His empty (MRC PPU Reagents and Services, DU 57719) were transformed into BL21 cells. Expression cultures were inoculated with 1% (v/v) of an overnight culture and were grown to OD_600_ of 0.3 at 37 °C, shaking at 180 rpm. The bacterial cultures were then cooled to 18 °C, induced at OD_600_ of 0.6-0.7 with 0.1 mM Isopropyl-β-D-thiogalactoside and incubated at 18 °C overnight. The following day, the cells were spun down at 4000 g for 25 min at 4 °C. Cell pellets were resuspended in lysis buffer containing 50 mM Tris-HCl pH 7.5, 150 mM NaCl, 5 mM MgCl_2_, 10% (v/v) glycerol, 0.5 mM TCEP (tris(2-carboxyethyl)phosphine), 10 mM Imidazole pH 8.0, 1 mM PMSF, and 1 mM benzamidine. Cells were lysed using an Avestin Emulsiflex apparatus at 20,000 psi at 4 °C. Lysate was clarified by centrifugation at 30,000 g for 30 min at 4 °C, and incubated with 200 μl of equilibrated nickel-NTA agarose (per 1 l culture) for 1 h rotating at 4 °C. The resin was washed twice with 50 mM Tris-HCl pH 7.5, 300 mM NaCl, 5 mM MgCl_2_, 10% glycerol, 0.5 mM TCEP, 20 mM Imidazole (high salt buffer), a total of 15 ml per 200 μl of resin, and twice with 50 mM Tris-HCl pH 7.5, 150 mM NaCl, 5 mM MgCl_2_, 10% glycerol, 0.5 mM TCEP, 20 mM Imidazole (low salt buffer), a total of 20 ml per 200 μl of resin. 1.5 mg of tissue lysate (prepared as described above) was diluted to 10 ml using low salt buffer to reduce Triton X-100 concentration to ∼0.03% and supplemented with GTP-γ-S at a final concentration of 100 μM. Tissue lysate was incubated with 200 μl nickel-NTA agarose with immobilized 6-His LRRK2 1-552 or 6-His for 2 h rotating at 4 °C. Resin was washed three times with 5 ml low salt buffer. An equal volume of elution buffer containing 500 mM Imidazole and low salt buffer was added to the resin and the slurry was transferred to Corning Costar spin-X centrifuge tube filters (Corning #8161). The resin was incubated with elution buffer for 20 min rotating at 4 °C. Protein was eluted in spin-X tubes by centrifugation at 500 g for 5 min at 4 °C. Protein was concentrated to 80 μl using Amicon Ultra centrifugal filters, 10K (#UFC901024). 15% of eluted protein was analyzed by immunoblotting.

### Microsome enrichment by fractionation

Fractionation to detect LRRK2 auto-phosphorylation at Ser1292 in mouse tissues was described previously [54]. Briefly, snap frozen mouse lungs were homogenized on ice in 15% (w/v) sedimentation buffer containing 3 mM Tris-HCl pH 7.4, 250 mM sucrose, 0.5 mM EGTA, 10 mM 2-glycerophosphate, 50 mM NaF, 5 mM sodium pyrophosphate, 1 μg/ml microcystin-LR, 1 mM sodium orthovanadate, and complete EDTA-free protease inhibitor cocktail (Roche). Homogenates were centrifuged at 3000 g for 10 min at 4 °C, and the supernatant was centrifuged again at 5000 g for 10 min at 4 °C to produce cleared homogenate. 200 μl cleared homogenate was supplemented with 10X lysis buffer (200 mM Tris-HCl pH 7.4, 1.5 M NaCl, 10 mM EGTA, 25 mM sodium pyrophosphate, 10 mM 2-glycerophosphate, 10 mM sodium orthovanadate, 1 μg/ml microcystin-LR, protease inhibitor cocktail tablet, and 10% Triton X-100) to a final concentration of 1X, and cleared homogenate was further supplemented with 0.1% (w/v) SDS and 0.5% (w/v) sodium deoxycholate. Cleared homogenate was incubated on ice for 1 h for complete lysis, and centrifuged at 17,000 g for 20 min at 4 °C. Supernatant from this centrifugation step was designated as the total fraction. The remainder of the cleared homogenate was centrifuged at 12,000 g for 10 min at 4 °C. The pellets were resuspended in 15% (v/v) sedimentation buffer supplemented with the 10X lysis buffer to a final concentration of 1X (as described above) to the volume of the original sample that was spun down. The resuspended pellets were termed crude mitochondrial fractions. The supernatant from the aforementioned centrifugation step was quantified using the Bradford method and 20 mg of supernatant was ultracentrifuged at 100,000 g for 1 h at 4 °C. The supernatant from this centrifugation step was designated the cytosol fraction. The pellets were washed twice with PBS and resuspended in 530 μl resuspension buffer (15% w/v sedimentation buffer supplemented with 10X lysis buffer). Resuspended pellets were sonicated three times for 15 sec and designated as microsomal fractions. 40 μg of the total fractions and the equivalent volumes of the remaining fractions were analyzed by immunoblotting.

### Cell Culture, Treatments, and Lysis

Mouse embryonic fibroblasts (MEFs) were cultured in Dulbecco’s modified eagle medium (DMEM) supplemented with 20% (v/v) fetal calf serum, 2 mM L-glutamine, 100 U/ml penicillin, 100 μg/ml streptomycin, non-essential amino acids and 1 mM sodium pyruvate. Primary lung fibroblasts were cultured in Dulbecco’s modified eagle medium: nutrient mixture F-12 (DMEM/F-12) supplemented with 20% (v/v) fetal calf serum, 2 mM L-glutamine, 100 U/ml penicillin, 100 μg/ml streptomycin, non-essential amino acids and 1 mM sodium pyruvate. Generation of A549 Rab29 knock-out cells was described previously [14], and A549 cells were cultured in DMEM supplemented with 10% (v/v) fetal calf serum, 2 mM L-glutamine, 100 U/ml penicillin and 100 μg/ml streptomycin. All cells were grown at 37 °C and 5% CO2 in a humidified atmosphere. Cell lines utilized for this study were tested regularly for mycoplasma contamination and confirmed as negative prior to experimental analysis. Cells were lysed in ice-cold lysis buffer containing 50 mM Tris/HCl pH 7.4, 1 mM EGTA, 10 mM 2-glycerophosphate, 50 mM sodium fluoride, 5 mM sodium pyrophosphate, 270 mM sucrose, supplemented with 1 μg/ml microcystin-LR, 1 mM sodium orthovanadate, complete EDTA-free protease inhibitor cocktail (Roche), and 1% (v/v) Triton X-100. Lysates were clarified by centrifugation for 15 min at 15,000 g at 4 °C. Protein concentrations of cell lysates were determined using the Bradford assay. MLi-2 inhibitor was dissolved in sterile DMSO, and treatment of cells with MLi-2 was for 90 minutes at a final concentration of 100 nM, unless otherwise indicated. Nigericin and monensin were dissolved in absolute ethanol and chloroquine diphosphate salt was dissolved in sterile water. Cell treatments were added at a dilution of 0.1% to cells. LLOMe was prepared in DMSO at a working concentration of 100 mM and cells were treated with a 1% dilution of stimulant. An equivalent volume of diluent was added to vehicle control cells where appropriate.

### Generation of mouse embryonic fibroblasts (MEFs)

Wildtype, heterozygous, and homozygous Rab29 knock-out or transgenic Rab29 overexpressing MEFs were isolated from littermate-matched mouse embryos at day E12.5, as described in a previous study [55]. The resulting embryo genotypes were produced following crosses between Rab29 knock-out/wildtype (heterozygous) mice or between transgenic Rab29/wildtype (heterozygous) mice. Rab29 knock-out and LRRK2[R1441C] mice were bred to create doubly modified Rab29 knock-out/LRRK2[R1441C], and Rab29-knock-out and VPS35[D620N] mice were bred to create doubly modified Rab29 knock-out/VPS35[D620N] mice. To generate MEFs for these mouse models, the following crosses between homozygous mice of the following genotypes were set up: Rab29 wildtype/LRRK2 or VPS35 wildtype, Rab29 knock-out/LRRK2 or VPS35 wildtype, Rab29 wildtype/LRRK2[R1441C] or VPS35[D620N], and Rab29 knock-out/LRRK2[R1441C] or VPS35[D620N]. The resulting embryos from each litter were of the same genotype and MEFs were isolated at day E12.5. Genotypes of all aforementioned mouse models were verified *via* allelic sequencing and immunoblot. Primary MEFs between passage 2 and 8 were used for all experimental analyses, except Figure 8 where MEFs ranged from passage 8 to 12.

**Figure 8.**
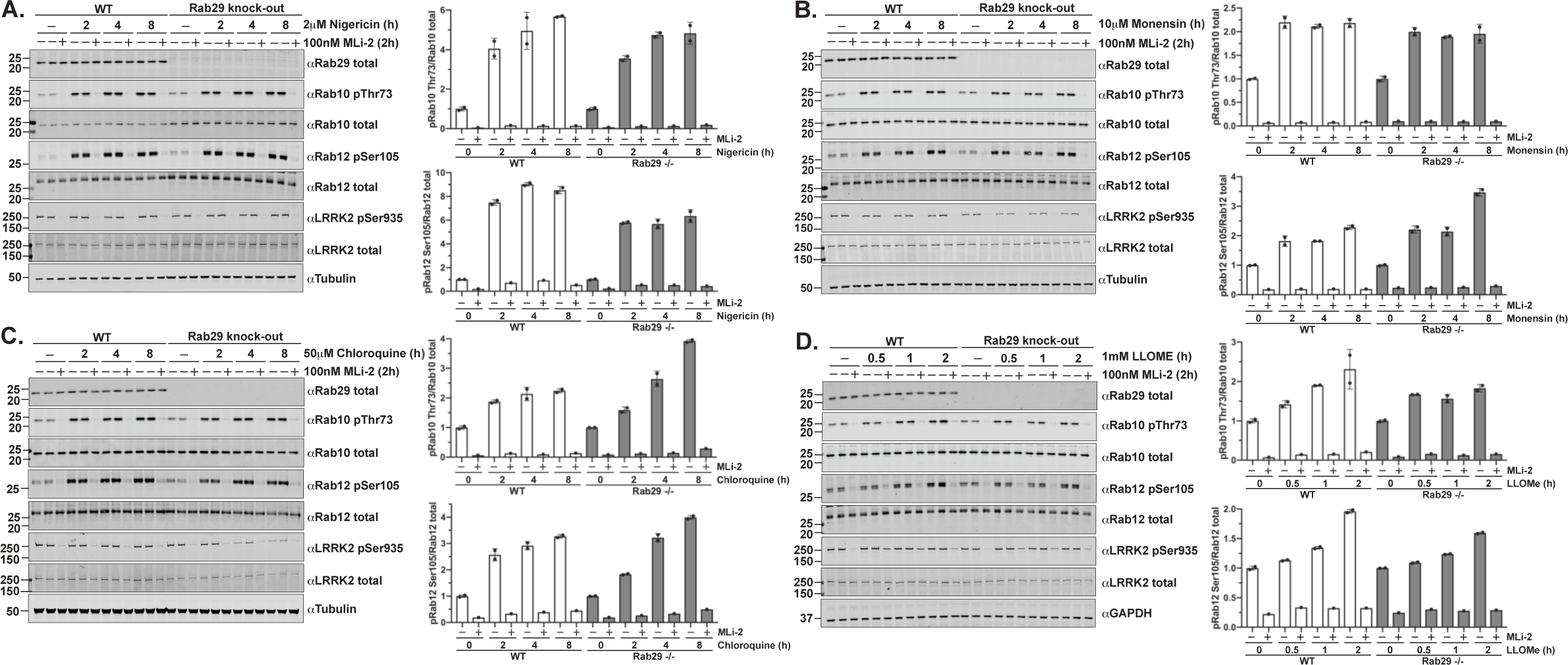
Cation ionophores nigericin and monensin, and lysosomal stressors chloroquine and LLOMe enhance LRRK2-mediated Rab10 and Rab12 phosphorylation in MEFs in a Rab29-independent manner. **(A-D)** Littermate matched wildtype and Rab29 knock-out MEFs were treated with vehicle or **(A)** 2 μM nigericin, **(B)** 10 μM monensin, **(C)** 50 μM chloroquine, or **(D)** 1 mM LLOMe for the indicated periods of time. Cells were treated with DMSO or 100 nM LRRK2 inhibitor MLi-2 for 2 hours prior to harvest. 15-20 μg of cell extract was subjected to quantitative immunoblot analysis with the indicated antibodies. The membranes were developed using the LI-COR Odyssey CLx Western Blot imaging system. Quantified data are presented as the ratios of phospho-Rab10/total Rab10 and phospho-Rab12/total Rab12, which were calculated using the Image Studio software. Values were normalized to the average of the vehicle-treated cells for each respective genotype. Similar results were obtained in two independent experiments for each agonist.

### siRNA-mediated knockdown of target proteins in MEFs

For siRNA knockdown of proteins of interest, ON-TARGETplus Mouse LRRK2 siRNA-SMARTpool (#L-049666-00-0005), ON-TARGETplus Mouse Rab32 siRNA-SMARTpool (#L-063539-01-0005) and ON-TARGETplus non-targeting pool (#D-001810-10-05) were purchased from Dharmacon. MEF cells were seeded in a 6-well format at 400,000 cells/well for transfection the following day. Cells were transfected using Lipofectamine RNAiMAX according to the manufacturer’s protocol. Briefly, 50 pmol siRNA was diluted in 150 μl opti-MEM and combined with 10 μl Lipofectamine RNAiMAX in 150 μl opti-MEM per well. The two mixtures were incubated together at room temperature for 5 min and 250 μl was added dropwise to cells, which were harvested 72 h after transfection.

### MLi-2 inhibitor washout in MEFs

Primary MEFs of the indicated genotypes in Figure 9 were seeded in a 6-well format for treatment the following day. At 50-60% confluence, MEFs were treated with vehicle (DMSO) or 100 nM MLi-2 for 48 h. To remove the MLi-2 inhibitor, cells were washed three times with warm, complete media. 15-20 min following the initial washes, cells were washed an additional two times with complete media. Cells were harvested with complete lysis buffer 30-360 min following the initial washes.

**Figure 9.**
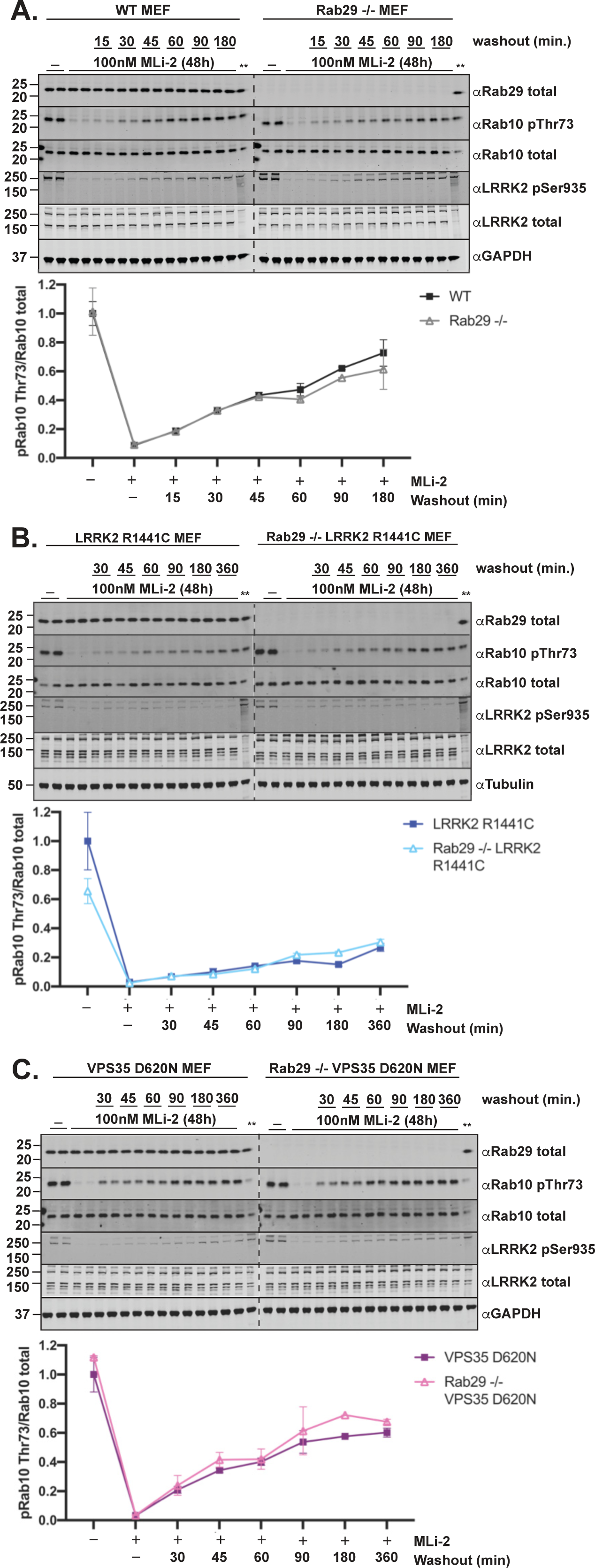
Rab29 knock-out does not impact recovery of LRRK2 activity following washout of LRRK2 inhibitor. **(A-C)** The indicated matched MEFs were treated with vehicle (DMSO) or 100 nM LRRK2 inhibitor MLi-2 for 48 h. The MLi-2 inhibitor was removed from cells through multiple washes with complete media for the indicated periods prior to cell lysis. 20 μg of whole cell extracts were subjected to quantitative immunoblot analysis with the indicated antibodies. The membranes were developed using the LI-COR Odyssey CLx Western Blot imaging system. Quantified data are presented as the ratios of phospho-Rab10/total Rab10, which were calculated using the Image Studio software. Values were normalized to the average of **(A)** wildtype MEFs treated with DMSO, **(B)** LRRK2[R1441C] MEFs treated with DMSO, or **(C)** VPS35[D620N] MEFs treated with DMSO. Cell extract derived from a wildtype MEF cell line (**) was added to each gel in order to accurately compare the Rab10 pThr73/Rab10 total ratios between genotypes in each panel. Data quantifications are presented as mean ± SD. Similar results were obtained in two separate experiments.

### Immunofluorescence microscopy

Littermate matched wildtype and transgenic Rab29 MEFs, and Rab29 knock-out MEFs were plated in a 6-well format on VWR 22×22mm cover slips (cat# 631-0125) that were soaked in absolute ethanol for 1 h prior to seeding 400,000 cells per well for each genotype. The following day, cells were fixed on coverslips with 4% paraformaldehyde in phosphate-buffered saline (PBS), pH 7.4, for 10 min. Cells were washed three times with 0.2% bovine serum albumin (BSA) diluted in PBS and permeabilized with 0.1% NP-40 diluted in PBS for 10 min. Cells were washed three times with 0.2% BSA and blocked with 1% BSA diluted in PBS for 1 h. Cells were washed three times with 0.2% BSA and incubated with primary antibodies against ACBD3 and Rab29 that detects mouse (MJF-30, clone 124), diluted to final concentrations of 10 μg/ml in 0.2% BSA, for 1 h. Cells were washed with 0.2% BSA three times for 5 min each and incubated with 1 μg/ml DAPI (Bisbenzimide Hoechst 33342 trihydrochloride, Sigma B2261) and secondary antibodies diluted 1:500 in 0.2% BSA for 1 h (Invitrogen donkey anti-rabbit IgG (H+L) Alexa Fluor 594 (cat# A-21207) and donkey anti-mouse IgG (H+L) Alexa Fluor 488 (cat# A21202)). Cells were washed three times with 0.2% BSA for 5 min each. The slides were rinsed in sterile water before mounting on VWR super premium microscope slides (cat# 631-0117) using Vectashield antifade mounting media (H-1000). All incubation steps were carried out at room temperature. Images were acquired using a Carl Zeiss LSM710 laser scanning confocal microscope (Carl Zeiss) using the 63X Plan-Apochromat objective (NA 1.4) and a pinhole chosen to provide a uniform optical section thickness in all fluorescence channels. The images were processed as a batch using ImageJ and brightness and contrast adjustments were kept constant for the same fluorescent channel.

### Generation of primary lung fibroblasts

Primary lung fibroblasts were derived from adult mice as previously described [56]. Briefly, lung tissue was harvested and transferred to a sterile tissue culture dish, washed twice with PBS and minced with a scalpel. Tissue fragments were transferred to a 75cm^2^ cell culture flask with suitable aeration and immersed in 10 ml DMEM/F12 supplemented with 2 mM L-glutamine, 100 U/ml penicillin, 100 μg/ml streptomycin, and 0.14 Wunsch units/ml collagenase (Liberase, Sigma-Aldrich 05401119001). The tissue fragments were digested with collagenase in a shaking incubator at 37 °C and 5% CO_2_ for 1 h. 20ml of DMEM/F12 supplemented with 20% (v/v) fetal calf serum, 2mM L-glutamine, 100 U/ml penicillin, and 100 μg/ml streptomycin was added, and the cell/tissue suspension was centrifuged at room temperature at 525 g for 5 min. The pellet was gently resuspended in fresh complete media and plated in a collagen-coated cell culture flask. To coat the cell culture flasks, Gibco Collagen I, rat tail (A10483-01) was diluted to a final concentration of 50 μg/ml in sterile-filtered 20 mM acetic acid. Tissue-culture flasks were coated with diluted collagen according to the manufacturer’s. Fibroblasts required ∼5 days to emerge from tissue fragments and regular media changes. Approximately 1.5 weeks following isolation, lung fibroblasts were passaged once, then seeded in 6-well plates for experimental analysis.

### Statistics

Graphs were made using Graphpad Prism 7 and 8 software. Error bars indicate SD. Two-tailed unpaired *t*-test or one-way ANOVA with Tukey’s multiple comparisons test were used to determine statistical significance. P-values < 0.05 were considered statistically significant. Based on previous publications that monitor Rab10 Thr73 phosphorylation and LRRK2 Ser935 phosphorylation in various genotypes, following vehicle or MLi-2 administration, [9,21,49,51,54], we determined that N = 4 per genotype for the studies outlined in Figures 6 and 7, gives > 80% power to reject the null hypothesis at α = 0.05.

## Results

### Expression of Rab29 in mouse tissues and cells

We raised two novel Rab29 monoclonal antibodies termed MJF-30-Clone-124 and MJF-30-Clone-104 that detect endogenous Rab29 in wildtype but not in Rab29 knock-out human A549 cells (Fig 1A). MJF-30-Clone-124 detected both mouse and human Rab29 whilst MJF-30-Clone-104 was human specific (Fig 1A). Immunoblotting of 6 mouse tissues (brain, spleen, lung, kidney, large intestine and spinal cord) and 3 primary mouse cell lines (mouse embryonic fibroblasts (MEFs), lung fibroblasts and bone marrow derived macrophages) that all express endogenous LRRK2, revealed that Rab29 is expressed ubiquitously but levels vary significantly between tissues and cells (Fig 1B). The highest expression is observed in macrophages and spleen, whilst low expression is seen in brain and spinal cord, and intermediate expression observed in other tissues and cells. LRRK2 was proteolyzed into at least two bands in many extracts but the expression of the two major upper bands was also lower in brain and spinal cord, and significantly higher in other tissues and cells. This is consistent with substantial evidence that LRRK2 and Rab29 are co-expressed in the same cells (MEFs, A549, macrophages, neutrophils) [14,15,43,57,58] (http://www.immprot.org/) as well as tissues (https://www.proteinatlas.org/).

### Rab29 knock-out does not impact basal LRRK2-mediated phosphorylation of Rab10 and Rab12 in mouse tissues and MEFs

To investigate the role of endogenous Rab29 in regulating LRRK2 pathway activity, we obtained Rab29 knock-out mice made available through the Wellcome Trust Sanger Institute, Infrafrontier EMMA mouse repository and the Michael J. Fox Foundation (see Methods). These mice have also been deployed in other studies mentioned above [43,44,46]. As a readout for LRRK2 pathway activity we measured phosphorylation of Rab10 at Thr73 [7], employing a well-characterized phospho-specific antibody [51]. Consistent with previous reports [44], the Rab29 knock-out mice were viable and displayed no overt phenotypes. We analyzed wildtype, heterozygous and Rab29 knock-out MEFs (Fig 2A) as well as wildtype and Rab29 knock-out lung (Fig 2B), spleen (Fig 2C), kidney (Fig 2D) and various brain sections (Fig S1) derived from littermate 6-month-old mice treated ± MLi-2 LRRK2 inhibitor (30 mg/kg, 2 h). Results from numerous independent experiments demonstrated that there was no significant difference in levels of LRRK2-phosphorylated Rab10, quantitated as a ratio with total Rab10, in wildtype or Rab29 knock-out MEFs, lung, spleen or kidney. In MEFs, we also analyzed LRRK2-mediated phosphorylation of Rab12 at Ser105 using a previously characterized phospho-specific antibody [21], which was also shown to be unaffected by Rab29 knock-out (Fig 2A). Rab29 knock-out also had no impact on LRRK2 expression or phosphorylation of LRRK2 at Ser935 (Fig 2, S1). As expected, MLi-2 treatment markedly reduced Rab10 phosphorylation levels in both wildtype and Rab29 knock-out mice and was accompanied by a decrease in LRRK2 Ser935 phosphorylation. We found that although MLi-2 administration reduced Ser935 phosphorylation in brain, the low basal levels of pRab10 observed were not further decreased (Fig S1), consistent with previous findings [51,59].

### Generation and characterization of Rab29 overexpressing transgenic mice

As the human genetic data point toward variants within the PARK16 locus enhancing expression of Rab29 [15,38,39], we generated a transgenic mouse strain in which Rab29 is constitutively overexpressed in all tissues. The mouse Rab29 cDNA with no epitope tags was knocked-into the ROSA26 locus, which is frequently employed to constitutively express proteins in mouse tissues [60] (Fig 3A). The heterozygous and homozygous Rab29 transgenic mice displayed no overt phenotype at 6 months of age, which were the oldest animals studied. Rab29 protein levels in brain, spleen, lung, kidney, large intestine and spinal cord derived from wildtype and homozygous transgenic Rab29 mice were analyzed at 3.5 and 6 months of age. This revealed that the highest overexpression of Rab29 was observed in brain (7-9-fold), kidney (5-7-fold) and large intestine (23-25-fold), with lower levels of overexpression observed in lung (<1.5-fold), spleen (<2-fold) and spinal cord (4-fold) (Fig 3B). The levels of other Rab proteins measured by immunoblotting (Rab8A, Rab10, Rab12, Rab32 and Rab35) were not impacted by overexpression of Rab29 (Fig 3B). There was no marked difference in the relative levels of Rab29 expression between 3.5 and 6-month-old mice (Fig 3B). We also analyzed Rab29 mRNA levels in wildtype, heterozygous and homozygous brain, lung and kidney from 3.5-month-old mice (Fig 3C). Consistent with protein levels, brain displayed the highest increase in Rab29 mRNA levels, namely 2-fold in heterozygous and ∼4-fold in homozygous Rab29 transgenic mice (Fig 3C). In transgenic homozygous kidney and lung, Rab29 mRNA levels were increased ∼3 and 2-fold, respectively (Fig 3C).

To demonstrate that the overexpressed Rab29 was functional, we undertook immunofluorescence studies of Rab29 knock-out and littermate matched wildtype and transgenic MEFs (Fig 4A). This revealed that the overexpressed transgenic Rab29 was correctly localized to the Golgi, similar to wildtype Rab29, and this was confirmed by co-staining with the ACBD3 Golgi marker. Consistent with our localization being specific, no Rab29 signal was observed in the Rab29 knock-out cells and enhanced levels of Rab29 were clearly observed in the transgenic cells (Fig 4A). Rab29 localization to the Golgi complex indicates that the protein is properly folded and active [30]. Rab29 binds to the N-terminus of LRRK2 and recent work suggests that the binding site is located within the first 552 residues of LRRK2 [41]. To obtain further evidence that the Rab29 expressed in the transgenic mouse was functional, we undertook affinity purification studies using a fragment of LRRK2 encompassing residues 1-552 in intestinal extracts that express the highest levels of Rab29. These studies confirmed that the transgenic Rab29 expressed in these extracts interacts with the LRRK2 N-terminal fragment, providing further evidence that the transgenic Rab29 is functionally competent (Fig 4B). Binding of endogenously expressed wildtype Rab29 to the LRRK2 N-terminal fragment was also observed in wildtype intestine extracts (Fig 4B).

### Transgenic overexpression of Rab29 does not impact basal LRRK2-mediated phosphorylation of Rab10 and Rab12 in mouse tissues and cells

We examined LRRK2-mediated Rab10 and Rab12 phosphorylation in wildtype and homozygous transgenic Rab29 mouse brain (Fig 5A), large intestine (Fig 5B), kidney (Fig 5C), spinal cord (Fig 5D), spleen (Fig 5E) and lung (Fig 5F) from 6-month-old mice treated for 2 h ± MLi-2 LRRK2 inhibitor (30 mg/kg). As expected, phosphorylation of Rab10 and Rab12 were robustly detected in all tissues and were significantly lowered by MLi-2 treatment (Fig 5). Notably, in the brain extracts, Rab12 phosphorylation was more sensitive to MLi-2 administration than Rab10 phosphorylation (Fig 5A). No significant increases in LRRK2-mediated phosphorylation of Rab10 or Rab12 were observed in any of the tissues of the Rab29 transgenic mice. Levels of LRRK2 or phosphorylation at Ser935 were also not impacted by Rab29 overexpression. Similar results were also observed in 3.5-month-old animals (Fig S2). We also studied MEFs and lung fibroblasts derived from wildtype and homozygous Rab29 transgenic mice in which Rab29 levels were increased ∼4-fold in both cell types, and here again observed no significant impact on LRRK2-mediated Rab10 or Rab12 phosphorylation (Fig S3). As expected, MLi-2 reduced Rab10 and Rab12 phosphorylation (Fig S3).

### Knock-out of Rab29 does not reduce elevated Rab10 and Rab12 phosphorylation in LRRK2[R1441C] knock-in mice

To study whether endogenous Rab29 was necessary for the previously reported elevated Rab10 phosphorylation observed in LRRK2[R1441C] knock-in MEFs and mouse tissues [26,51], we generated LRRK2[R1441C] knock-in MEFs as well as 6-month-old LRRK2[R1441C] ± Rab29 knock-out mice. As a control, we also generated matched LRRK2 wildtype ± Rab29 knock-out MEFs for comparison. In MEFs, as reported previously [51], the LRRK2[R1441C] knock-in mutation enhanced Rab10 phosphorylation around 3-fold compared to wildtype (Fig 6A). Knock-out of Rab29 had no impact on Rab10 phosphorylation (Fig 6A). As reported previously [61], the R1441C mutation reduced Ser935 phosphorylation, which was also not impacted by Rab29 knock-out (Fig 6A). In mouse brain (Fig 6B), lung (Fig 6C), kidney (Fig 6D), spleen (Fig 6E), large intestine (Fig S4A) and spinal cord (Fig S4B), we also observed that knock-out of Rab29 in LRRK2[R1441C] knock-in mice had no significant impact on Rab10 phosphorylation. The LRRK2[R1441C] mutation enhanced Rab10 phosphorylation between 1.5 to 2-fold in lung (Fig 6B), kidney (Fig 6C), spleen (Fig 6D), and large intestine (Fig S4A). No significant increase in Rab10 phosphorylation was observed in the brain (Fig 6B) and spinal cord (Fig S4B) of LRRK2[R1441C] knock-in mice. However, moderate sensitivity to MLi-2 was observed in the LRRK2[R1441C] knock-in mice, which was less apparent in wildtype animals (Fig 6B). Interestingly, pRab12 in the brain samples was more clearly regulated by LRRK2[R1441C] compared to Rab10 (Fig 6B). MLi-2 markedly decreased pRab12 levels 2.6-fold in the LRRK2[R1441C] and more modestly in wildtype brain samples, around 1.3-fold (Fig 6B). We observed ∼2-fold increase in pRab12 levels in LRRK2[R1441C] brain samples compared to wildtype (Fig 6B). In contrast, phosphorylation of Rab10 was not significantly impacted by the LRRK2[R1441C] mutation or by MLi-2 in the same extracts (Fig 6B). Similar results were also observed in the lung (Fig 6C), kidney (Fig 6D) and spleen (Fig 6E). Moreover, pRab10 or pRab12 levels were not significantly impacted by knock-out of Rab29 in these tissues.

We also immunoblotted for the pRab specific phosphatase PPM1H [33], and found that this was most highly expressed in the brain tissue, but found that its levels were not impacted by loss of Rab29 in multiple tissues (Fig 6B, 6C, 6D and 6E). In addition, we immunoblotted kidney tissue for the mannose-6-phosphate receptor (M6PR) that had previously been reported to be more highly expressed in Rab29 knock-out mice [46]. We confirm this finding and found that M6PR was expressed at ∼2.5 fold higher levels in the Rab29 knock-out kidney, but expression was not significantly affected by the LRRK2[R1441C] mutation or MLi-2 administration (Fig 6D). Mannose-6-phosphate receptor is often overexpressed when its trafficking is compromised.

We also attempted to measure endogenous phosphorylation of LRRK2 at Ser1292, an autophosphorylation site, using a commercially available antibody [10,54]. This site is challenging to detect, as stoichiometry of phosphorylation at this site is believed to be low. In addition to attempting to measure endogenous LRRK2 Ser1292 phosphorylation in total extracts, we also purified microsomal enriched fractions from lungs (that express highest levels of LRRK2), a method that was reported to facilitate detection of endogenous Ser1292 [54]. We were unable to robustly detect and quantify LRRK2 Ser1292 phosphorylation in either the total or microsome enriched fractions under conditions in which pRab10 levels were strongly detected (Fig S5).

### Knock-out of Rab29 does not reduce elevated Rab10 and Rab12 phosphorylation in VPS35[D620N] knock-in mice

To investigate whether endogenous Rab29 was necessary for the elevated Rab10 phosphorylation observed in VPS35[D620N] MEFs and mouse tissues [21], we generated VPS35[D620N] knock-in MEFs and 6-month-old VPS35[D620N] ± Rab29 knock-out mice. As a control, we also generated matched VPS35 wildtype ± Rab29 knock-out animals. As reported previously [21], VPS35[D620N] knock-in mutation enhanced Rab10 and Rab12 phosphorylation to a greater extent than is observed with the LRRK2[R1441C] pathogenic mutation (compare Fig 6 with Fig 7). Knock-out of Rab29 had no significant impact on the elevated Rab10 phosphorylation in the VPS35[D620N] knock-in MEFs (Fig 7A) or in mouse brain (Fig 7B), lung (Fig 7C), kidney (Fig 7D), spleen (Fig 7E), large intestine (Fig S6A), and spinal cord (Fig S6B). In brain extracts derived from VPS35[D620N] mice, a moderate 1.5-fold increase in Rab10 phosphorylation was observed, which decreased with MLi-2 administration, but was also not significantly impacted by Rab29 knock-out (Fig 7B). In contrast, the VPS35[D620N] mutation increased Rab12 phosphorylation ∼3-fold in brain extracts, which was suppressed by MLi-2 administration (Fig 7B). The VPS35[D620N] mutation also enhanced Rab10 and Rab12 phosphorylation in the lung (Fig 7C), kidney (Fig 7D) and spleen (Fig 7E). Here again, knock-out of Rab29 had no impact on Rab10 or Rab12 phosphorylation in any tissue studied. Consistent with previous results [21], the phosphorylation of Ser935 was not impacted by the VPS35[D620N] mutation. Knock-out of Rab29 had no impact on Ser935 phosphorylation in these MEFs and mouse tissues (Fig 7). We also found that M6PR was expressed at 2.5-fold higher levels in the Rab29 knock-out kidney and are increased ∼1.7-fold in the VPS35[D620N] mice (Fig 7D).

### Monovalent cation ionophore antibiotics nigericin and monensin stimulate LRRK2-mediated phosphorylation of Rab10 and Rab12 in a Rab29 independent manner

We next investigated whether we could identify agonists that stimulate LRRK2 pathway activity. We profiled a panel of agonists and stressors, which led to the finding that structurally related antibiotics nigericin and monensin markedly enhanced phosphorylation of Rab10 and Rab12 in wildtype MEFs (Fig 8A and 8B). This was blocked by treatment with the inhibitor MLi-2, indicating that nigericin and monensin stimulated Rab10 and Rab12 phosphorylation *via* LRRK2. Nigericin is derived from *Streptomyces hygroscopicus* [62], and monensin from *Streptomyces cinnamonensis* [63]. These agents function as monovalent cation ionophores inducing pleiotropic effects on vesicle trafficking responses [63]. We found that treatment of MEFs with 2 μM nigericin, over a 2 to 8 h time course, enhanced Rab10 phosphorylation 3.5 to ∼6-fold and Rab12 phosphorylation ∼6 to 9-fold (Fig 8A). 10 μM monensin over this period enhanced Rab10 phosphorylation up to ∼2.3-fold and Rab12 phosphorylation up to ∼3.5-fold in MEFs (Fig 8B). Knock-out of Rab29 had no significant impact on the stimulation of Rab10 and Rab12 phosphorylation observed with nigericin or monensin (Fig 8A, 8B). We also found that nigericin and monensin significantly stimulated Rab10 phosphorylation in A549 cells in a manner that was also unaffected by CRISPR knock-out of Rab29 (Fig S7).

Recent work has reported that lysosomotropic agents including chloroquine [64] and the peptide LLOMe (L-leucyl-L-leucine methyl ester) [65] enhance Rab10 phosphorylation [57,66–68]. We confirmed that both 50 μM chloroquine and 1 mM LLOMe (concentrations used in previous studies) enhance Rab10 as well as Rab12 phosphorylation in manner that was suppressed by MLi-2 (Fig 8C, 8D). The enhancement of Rab10 and Rab12 phosphorylation by chloroquine over an 8-hour time period was up to ∼3.5 and 4-fold, respectively (Fig 8C). The increase in Rab10 and Rab12 phosphorylation upon LLOMe stimulation was lower than observed with nigericin, monensin, and chloroquine, approaching ∼2-fold in MEFs (Fig 8D). Knock-out of Rab29 had no significant effect on Rab10 or Rab12 phosphorylation induced by chloroquine or LLOMe (Fig 8C, 8D). For the 1 mM LLOMe stimulation, we limited our analysis to 2 h, because MEF cells started to detach from tissue culture plates by 4 h.

### Rate of recovery of Rab10 phosphorylation after washout of MLi-2 LRRK2 inhibitor is not impacted by Rab29 knock-out

We next investigated whether endogenous Rab29 affected the rate at which Rab10 was re-phosphorylated following washout of MLi-2 in MEFs. We treated wildtype (Fig 9A), LRRK2[R1441C] (Fig 9B) or VPS35[D620N] (Fig 9C) ± Rab29 knock-out MEFs with MLi-2 to reduce pRab10 to undetectable levels (100 nM, 48 h). MLi-2 was removed by sequential changes of medium over a 15 min period, and phosphorylation of Rab10 quantified at time points up to 6 h. Under these conditions, we observed that the rate of recovery in the 3 cell lines was not affected by Rab29 knock-out. For the wildtype and VPS35[D620N] MEFs we observed 60-70% recovery of Rab10 phosphorylation within 6 h. However, in the LRRK2[R1441C] MEFs, recovery of pRab10 was only ∼30% after 6 h, and it is possible that the higher affinity of MLi-2 for this pathogenic mutant might account for this.

### Evidence that Rab32 is not compensating for loss of Rab29 in regulating LRRK2

Finally, we explored whether Rab32, which is closely related to Rab29 and reported to interact with LRRK2 [41,69], could contribute to regulation of LRRK2 activity in Rab29 knock-out MEFs. siRNA knockdown reduced Rab32 expression by 70-80%, however this had no impact on Rab10 phosphorylation in Rab29 knock-out wildtype LRRK2 or LRRK2[R1441C] knock-in cells (Fig 10). Knockdown of LRRK2 in parallel experiments, as expected, markedly reduced Rab10 phosphorylation (Fig 10). By immunoblotting analysis of wildtype and Rab29 knock-out MEFs we were unable to detect Rab38, which is also related to Rab29 and Rab32. This is consistent with previous high-resolution proteomic analysis of MEFs in which Rab38 was not detected [7].

**Figure 10.**
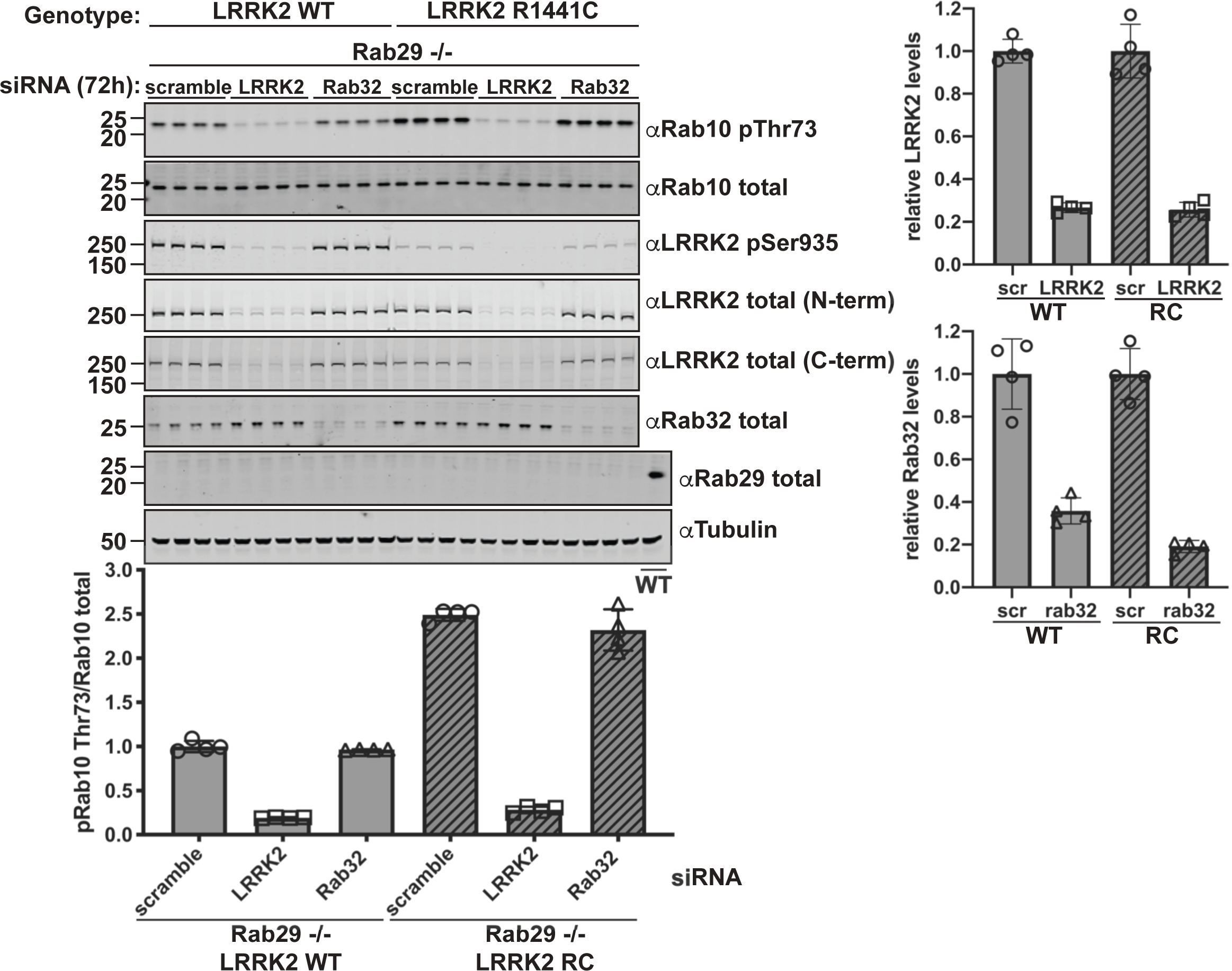
Knockdown of Rab32 does not impact LRRK2 activity in the absence of Rab29. The indicated matched primary MEFs were transfected with Dharmacon smartPOOL siRNA targeting either LRRK2, Rab32, or control scrambled non-targeting siRNA. The cells were lysed 72 h post-transfection. 20 μg of whole cell extracts were subjected to quantitative immunoblot analysis with the indicated antibodies. The membranes were developed using the LI-COR Odyssey CLx Western Blot imaging system. Quantified data are presented as the ratios of phospho-Rab10/Total Rab10 and were calculated using the Image Studio software. Values were normalized to the average of Rab29 knock-out MEFs treated with scrambled siRNA. The ratio of LRRK2 or Rab32 expression divided by the loading control was used to determine siRNA knockdown efficiency, and values were normalized to the average of the scrambled siRNA treated MEFs. Data quantifications are represented as mean ± SD. Similar results were obtained in three separate experiments.

## Discussion

As outlined in the introduction, considerable evidence supports the view that transient overexpression of Rab29 recruits the bulk of cellular LRRK2 to the Golgi surface, leading to its activation. In overexpression studies, the LRRK2[R1441C/G] mutants are more readily activated by Rab29, which could explain why this mutation elevates LRRK2 activity. However, our results suggest that knock-out of endogenous Rab29 has no significant impact on endogenous LRRK2 activity, assessed by monitoring pRab10 and pRab12 levels in six mouse tissues as well as MEFs and lung derived fibroblasts. Moreover, we also found that knock-out of Rab29 does not impact elevated Rab10 or Rab12 phosphorylation observed in the LRRK2[R1441C] knock-in MEFs or mouse tissues. This would suggest that endogenous Rab29 is not sufficient to explain the elevated activity of the LRRK2[R1441C] pathogenic mutant. We also found that Rab29 knock-out had no effect on the elevated LRRK2-mediated Rab10 or Rab12 phosphorylation observed in VPS35[D620N] knock-in MEFs and mouse tissues. To our knowledge, there is no evidence implicating Rab29 in mediating the effects of the VPS35[D620N] mutation. We also find that in brain extracts, Rab12 phosphorylation appears to be more robustly impacted by LRRK2 inhibitors and pathogenic mutations than Rab10 phosphorylation, consistent with a recent study [70].

In this study, we also report that nigericin and monensin markedly enhance Rab10 and Rab12 phosphorylation in MEFs and A549 cells, in a manner that is blocked by LRRK2 inhibitors. Nigericin and monensin function as broad monovalent cation ionophores, and have been reported to have pleiotropic effects on vesicular trafficking pathways [63]. We have also confirmed recent reports that lysosomotropic agents including chloroquine and LLOMe enhance Rab10 phosphorylation [57,66–68], and find that these agents also promote Rab12 phosphorylation (Fig 8A and 8B). We find that in MEFs, monensin, chloroquine and LLOMe induce pRab10 and pRab12 around 2 to 4-fold, compared to 4 to 9-fold effects observed with nigericin. A recent report suggested that chloroquine induced phosphorylation of Rab10 in RAW264.7 macrophages in a manner that was largely blocked by siRNA knock-down of Rab29 [67]. In contrast, we find that in MEFs, knock-out of Rab29 does not impact moderate Rab10 or Rab12 phosphorylation induced by chloroquine as well as the other agents we have tested (nigericin, monensin, LLOMe). It should be noted that Rab29 is highly expressed in macrophages [47] and in future work, it would be important to utilize Rab29 knock-out models to verify whether Rab29 plays a role in regulating LRRK2 pathway activity through agents such as chloroquine in primary macrophages. Further work is required to understand the mechanism by which nigericin, monensin and lysosomotropic agents promote LRRK2-mediated Rab protein phosphorylation. It would be interesting to explore whether these agonists promote recruitment of LRRK2 to particular membrane compartments, thereby activating LRRK2 and promoting Rab10 and Rab12 phosphorylation at that location. Recruitment of LRRK2 to membranes has been proposed previously to be a key mechanism by which LRRK2 activity is regulated [29,57,68].

It is possible that other Rab proteins, or even other regulators that have not yet been characterized, could also interact with and activate LRRK2 in a similar manner to Rab29 by recruiting LRRK2 to a variety of cellular membranes. In the absence of Rab29 overexpression, LRRK2 is widely distributed in cells with ∼90% cytosolic localization and ∼10% localization on a variety of cellular membranes [14,57,71,72]. A small fraction of LRRK2 appears to be localized in the Golgi region without Rab29 overexpression [14–17,73], which may explain why Rab29 knock-out does not have a noticeable impact on Rab10 of Rab12 protein phosphorylation when measured in a whole cell or tissue extract. In future work, it will be important to develop assays in which the pool of endogenous LRRK2 that resides at the Golgi could be specifically assessed. Since LRRK2 is likely in equilibrium between membranes and the cytosol [29], such interactions may be transient and more challenging to capture quantitatively. Indeed, by elevating the local concentration of Rab29 on the Golgi by exogenous expression, this pool was more readily detected. It will be important to define whether the activity and localization of the endogenous pool of Golgi-resident LRRK2 is dependent upon Rab29. It will also be necessary to further study whether endogenous LRRK2 located on other specific membrane compartments relies on other Rab proteins or regulators for this localization, and whether this contributes to the total cellular LRRK2 activity measured in cell extracts. It would also be important to define more precisely the residues in LRRK2 that bind Rab29 and investigate how subtle mutations that prevent Rab29 binding, impact cellular LRRK2 activity. Whether other Rab proteins bind to the same or different sites in LRRK2 should also be investigated.

The ability of exogenous Rab29 to recruit LRRK2 to the Golgi (or other compartments to which it is targeted) confirms the ability of Rab29 to bind LRRK2 in cells. Yet our knock-in and knock-out models failed to reveal clues regarding the functional significance of this interaction. Rab29 is a relatively poorly abundant Rab and may play a specific role in macrophages and dendritic cells where it is most abundant. Indeed, LRRK2 activation triggered by Rab29 could occur in a specific cell type or tissue, or following a physiological stimulus, stress or infection that we have not investigated. Our data do not rule out the possibility that Rab29 knock-out is compensated by another cellular protein(s) other than Rab32. However, our findings with transgenic mice that overexpress Rab29 from 1.5 to 25-fold, without enhancing LRRK2-mediated Rab10 or Rab12 phosphorylation, reveal that increasing Rab29 expression is not sufficient to stimulate the activity of endogenous LRRK2.

We have previously shown that Rab3, Rab8A, Rab10 Rab12, Rab35 and Rab43 are the main substrates of LRRK2 [7,26,74], and have herein monitored changes in phosphorylation of Rab10 and Rab12. We therefore cannot exclude the possibility that Rab29 preferentially activates phosphorylation of the Rab proteins we have not assayed, but we consider this unlikely because this set of Rab proteins appear to be coordinately phosphorylated in all cultured cell experiments that we have carried out to date [26,33,51]. It has also been suggested that screens could be undertaken to identify potentially therapeutic compounds that block Rab29 binding to LRRK2, however our data suggest that such agents may not be effective at reducing basal LRRK2 activity, unless these chemicals also block LRRK2 binding to other regulators transport it to cellular membranes. Finally, we propose that the new Rab29 monoclonal antibodies we have developed could be exploited to better understand how Parkinson’s mutations within the PARK16 locus impact Rab29 protein expression.

## Acknowledgements

We thank Suzanne Pfeffer (Stanford), Shalini Padmanabhan (The Michael J. Fox Foundation for Parkinson’s Research), and Sven M. Lange (MRC PPU) for helpful discussions, Gail Gilmour and Shauna Channon for mouse genotyping, and the excellent technical support of the MRC PPU, including the MRC PPU tissue culture team (coordinated by Edwin Allen) and MRC PPU Reagents and Services antibody teams (coordinated by Hilary McLauchlan and James Hastie).

## Author Contributions

A.F.K. designed, executed all experiments in this study apart from Figure 1A and Figure S7, analyzed and interpreted data and wrote the manuscript with D.R.A. J.B.F. discovered that nigericin and monensin stimulated LRRK2-mediated Rab10 phosphorylation and undertook experiments shown in Figure S7. P.L. designed and executed experiments that led to the development and characterization of the Rab29 MJF-30 Clone-104 and Clone-124 antibodies as well as undertaking work shown in Figure 1A. E.G.V suggested the Rab29 affinity purification study with the LRRK2 1-552 fragment, provided the expression construct for this study and assisted with the analysis and interpretation of data. N.K.P. oversaw and coordinated the generation of the Rab29 transgenic mice as well as the Rab29 MJF-30 antibodies. D.R.A. assisted with experimental design, analysis and interpretation of data and wrote the paper with A.F.K.

## Reagents and Data availability

In the materials and methods section, we have stated where all reagents used in this study can be obtained. All MRC-PPU reagents are available from our Reagents and Services division (https://mrcppureagents.dundee.ac.uk/). All primary data can be obtained by emailing DRA.

## Supplementary Figures

**Supplementary Figure 1.**
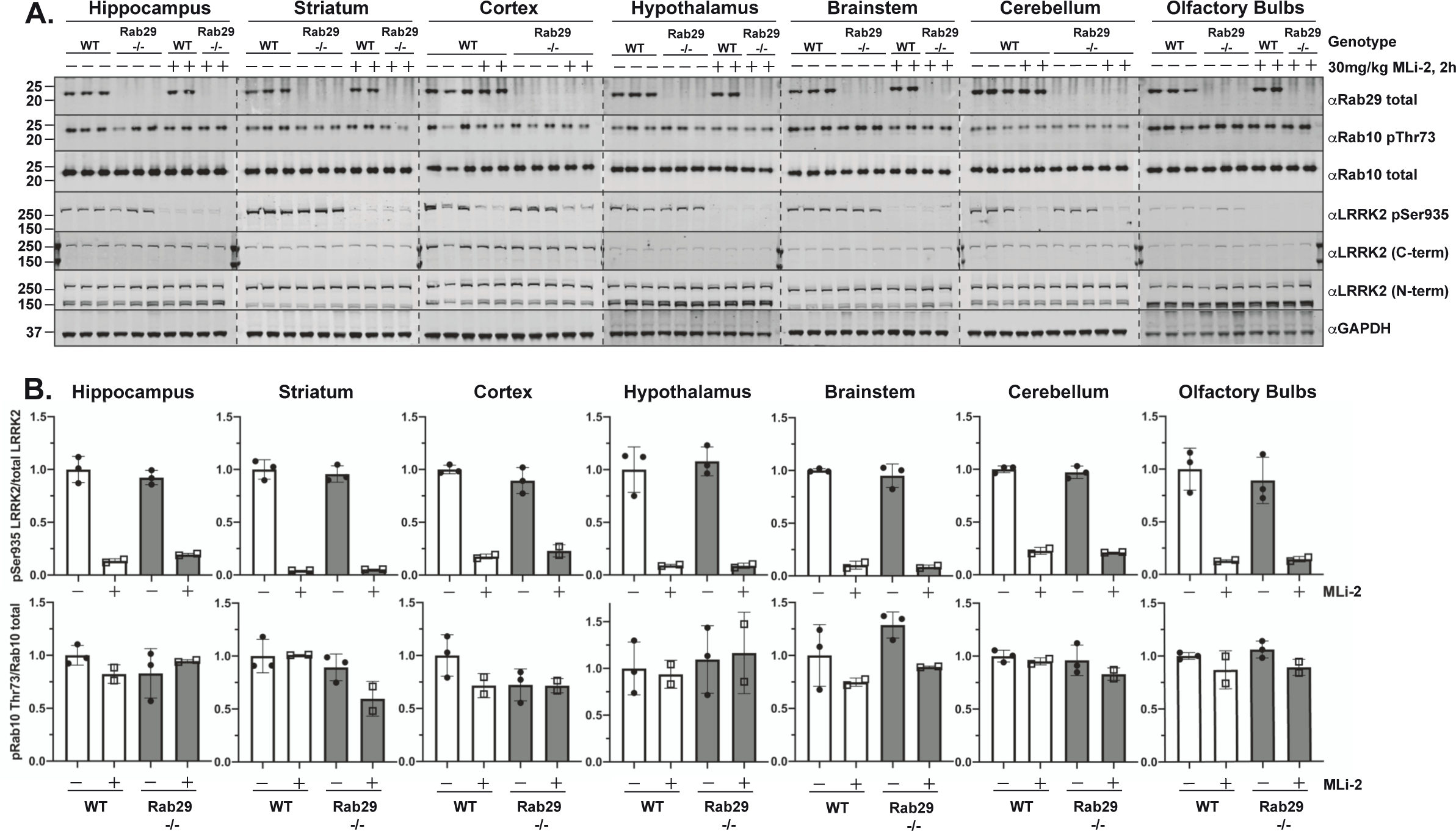
Knock-out of Rab29 does not affect LRRK2 Ser935 phosphorylation or LRRK2-mediated Rab10 phosphorylation in brain. **(A**) As in Figure 2 (B-D), 6-month-old, littermate-matched wildtype (WT) and Rab29 knock-out (-/-) mice were administered with vehicle (40% (w/v) (2-hydroxypropyl)-β-cyclodextrin) or 30 mg/kg MLi-2 dissolved in vehicle by subcutaneous injection 2 h prior to tissue collection. 40 μg of tissue extracts derived from 7 different brain sections were analyzed by quantitative immunoblot with the indicated antibodies. Each lane represents tissue extract derived from a different mouse. **(B)** Quantified data are presented as the mean ± SD of phospho-Rab10/Total Rab10 ratios and phospho-LRRK2/total LRRK2 ratios, and values were quantified using the Image Studio software.

**Supplementary Figure 2.**
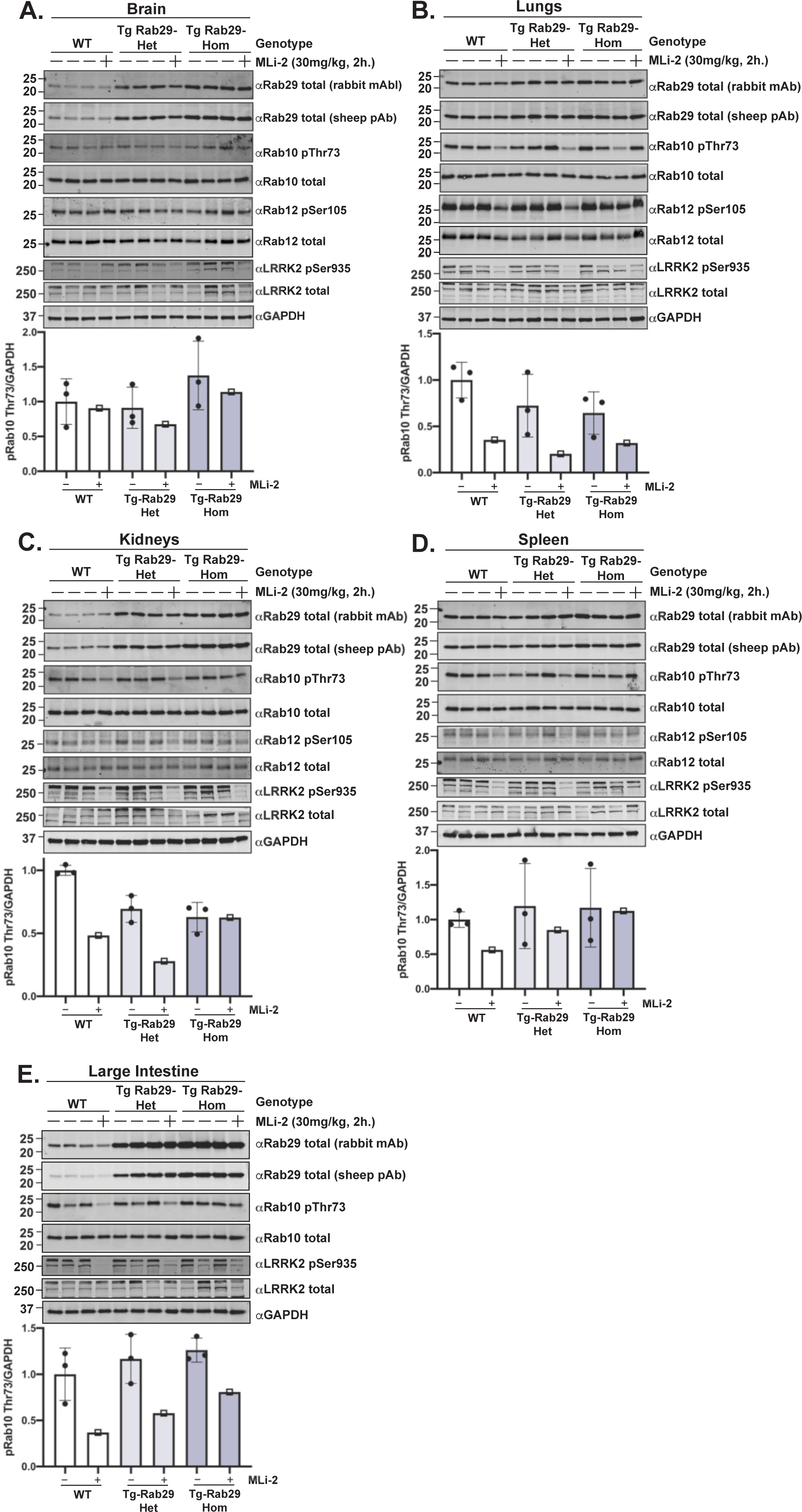
Overexpression of Rab29 in tissues from 3.5-month-old mice does not impact LRRK2-mediated Rab10 phosphorylation. **(A-E)** 3.5-month-old wildtype (WT), heterozygous (Tg Rab29 Het), and homozygous (Tg Rab29 Hom), transgenic Rab29 overexpressing mice were administered with vehicle (40% (w/v) (2-hydroxypropyl)-β-cyclodextrin) or 30 mg/kg MLi-2 dissolved in vehicle by subcutaneous injection 2 h prior to tissue collection. 40 μg of tissue extracts were subjected to quantitative immunoblot analysis with the indicated antibodies. The membranes were developed using the LI-COR Odyssey CLx Western Blot imaging system. Quantified data are presented as the phospho-Rab10/GAPDH ratios calculated using the Image Studio software. Values were normalized to the average of the wildtype, vehicle treated mice. Each lane represents a tissue sample derived from a different animal. Quantifications are presented as mean ± SD.

**Supplementary Figure 3.**
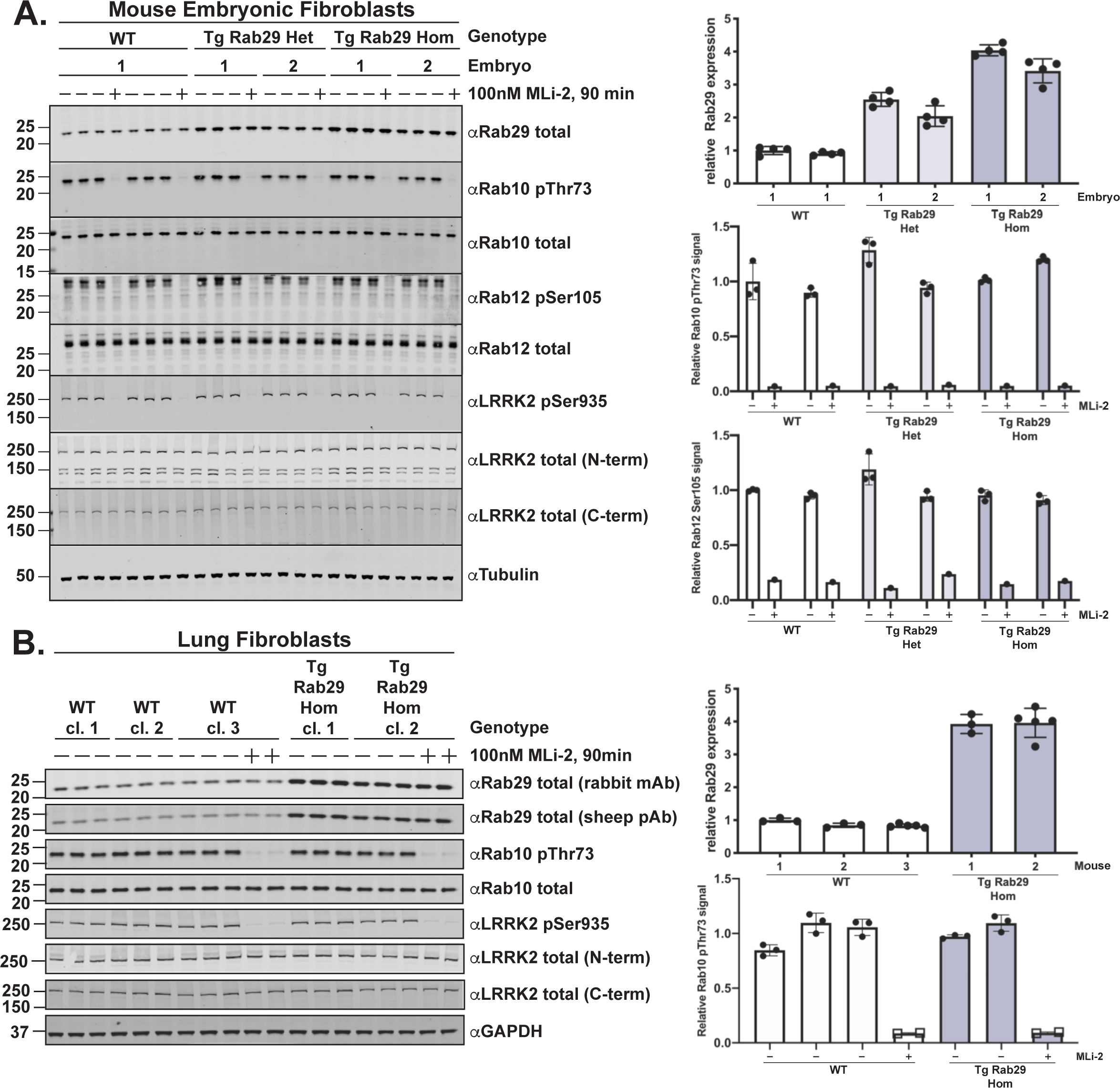
Overexpression of Rab29 in MEFs or primary lung fibroblasts does not impact LRRK2-mediated Rab10 phosphorylation. **(A)** Littermate-matched wildtype (WT), heterozygous (Tg Rab29 Het), or homozygous (Tg Rab29 Hom), transgenic Rab29 overexpressing MEFs were treated with vehicle (DMSO) or 100 nM LRRK2 inhibitor MLi-2 for 90 min prior to harvest. 20 μg of whole cell extracts were subjected to quantitative immunoblot analysis with the indicated antibodies. Technical replicates represent cell extract obtained from a different dish of cells. The membranes were developed using the LI-COR Odyssey CLx Western Blot imaging system. Quantified data are presented as the mean ± SD of phospho-Rab10/total Rab10 and phospho-Rab12/total Rab12. Rab29 levels were quantified by calculating the ratio of total Rab29/GAPDH. Data quantifications were undertaken using the Image Studio software and values were normalized to the average of wildtype MEFs treated with DMSO. **(B)** Primary lung fibroblasts derived from 3 different wildtype (WT) mice and 2 different transgenic, homozygous Rab29 overexpressing (Tg Rab29 Hom) mice were treated with vehicle (DMSO) or 100 nM LRRK2 inhibitor MLi-2 for 90 min prior to harvest. 20 μg of whole cell extracts were subjected to quantitative immunoblot analysis with the indicated antibodies. Technical replicates represent cell extract obtained from a different dish of cells. Quantified data are presented as the mean ± SD of phospho-Rab10/total Rab10. Rab29 levels were quantified by the ratio of total Rab29/GAPDH. Data quantifications were undertaken using the Image Studio software and values were normalized to the average of wildtype primary lung fibroblasts treated with DMSO.

**Supplementary Figure 4.**
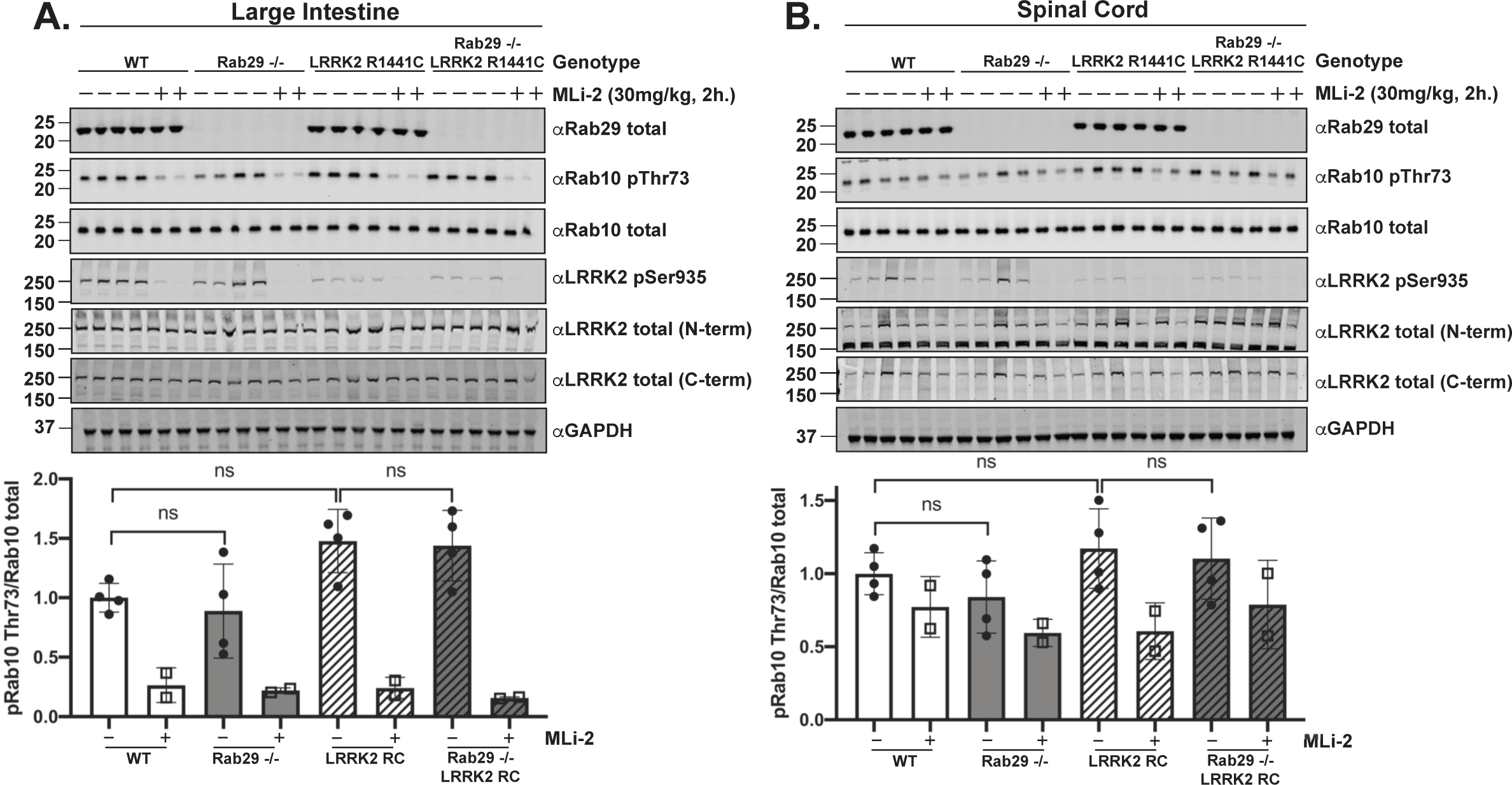
Knock-out of Rab29 does not reduce elevated Rab10 phosphorylation in pathogenic LRRK2[R1441C] knock-in mice. **(A-B)** As in Figure 5 (B-D), the indicated 6-month-old matched mice were administered with vehicle (40% (w/v) (2-hydroxypropyl)-β-cyclodextrin) or 30 mg/kg MLi-2 dissolved in vehicle by subcutaneous injection 2 h prior to tissue collection. 40 μg of tissue extracts were subjected to quantitative immunoblot analysis with the indicated antibodies. The membranes were developed using the LI-COR Odyssey CLx Western Blot imaging system. Quantified data are presented as the ratios of phospho-Rab10/total Rab10, calculated with the Image Studio software. Quantifications are presented as mean ± SD, normalized to vehicle treated, wildtype animals. Each lane represents a tissue sample from a different animal. Data were analyzed by one-way ANOVA with Tukey’s multiple comparisons test and no statistical significance was determined between the genotypes. Wildtype vs LRRK2[R1441C]: P = 0.1413 (A), P = 0.7459 (B). Wildtype vs Rab29 knock-out: P = 0.9455 (A), P = 0.7867 (B). LRRK2[R1441C] vs Rab29 knock-out LRRK2[R1441C]: P = 0.9973 (A), P = 0.9768 (B).

**Supplementary Figure 5.**
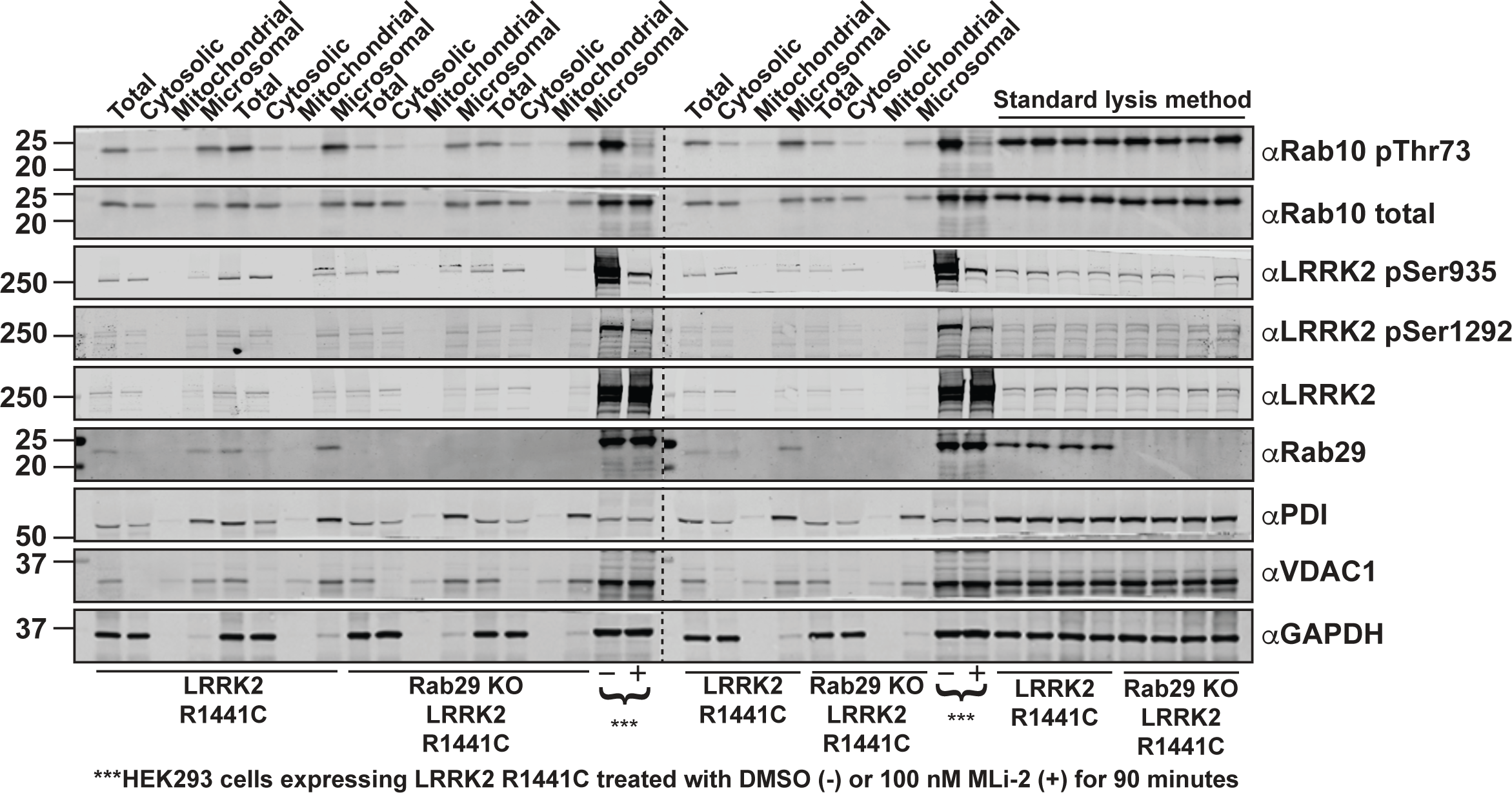
Microsomal enrichment by fractionation of LRRK2[R1441C] knock-in and Rab29 knock-out LRRK2[R1441C] knock-in lungs. Lung tissues from mice of the indicated genotypes were subjected to enrichment by fractionation to produce total, cytosolic, crude mitochondrial and microsomal fractions. 40 μg of the total fractions and the equivalent volumes of the remaining fractions were subjected to quantitative immunoblot analysis with the indicated antibodies. Antibodies against PDI, VDAC1, and GAPDH were used as markers for microsomal, mitochondrial and cytosolic fractions, respectively. HEK293 cells expressing FLAG-LRRK2 R1441C treated with or without 100 nM MLi-2 for 90 minutes were run in parallel as a control to confirm that phosphorylation of Ser1292 is LRRK2-mediated. 40 μg of lung tissue lysates from LRRK2[R1441C] and Rab29 knock-out LRRK2[R1441C] mice that were processed using the standard lysis method outlined in this study, were also run in parallel. The membranes were developed using the LI-COR Odyssey CLx Western Blot imaging system.

**Supplementary Figure 6.**
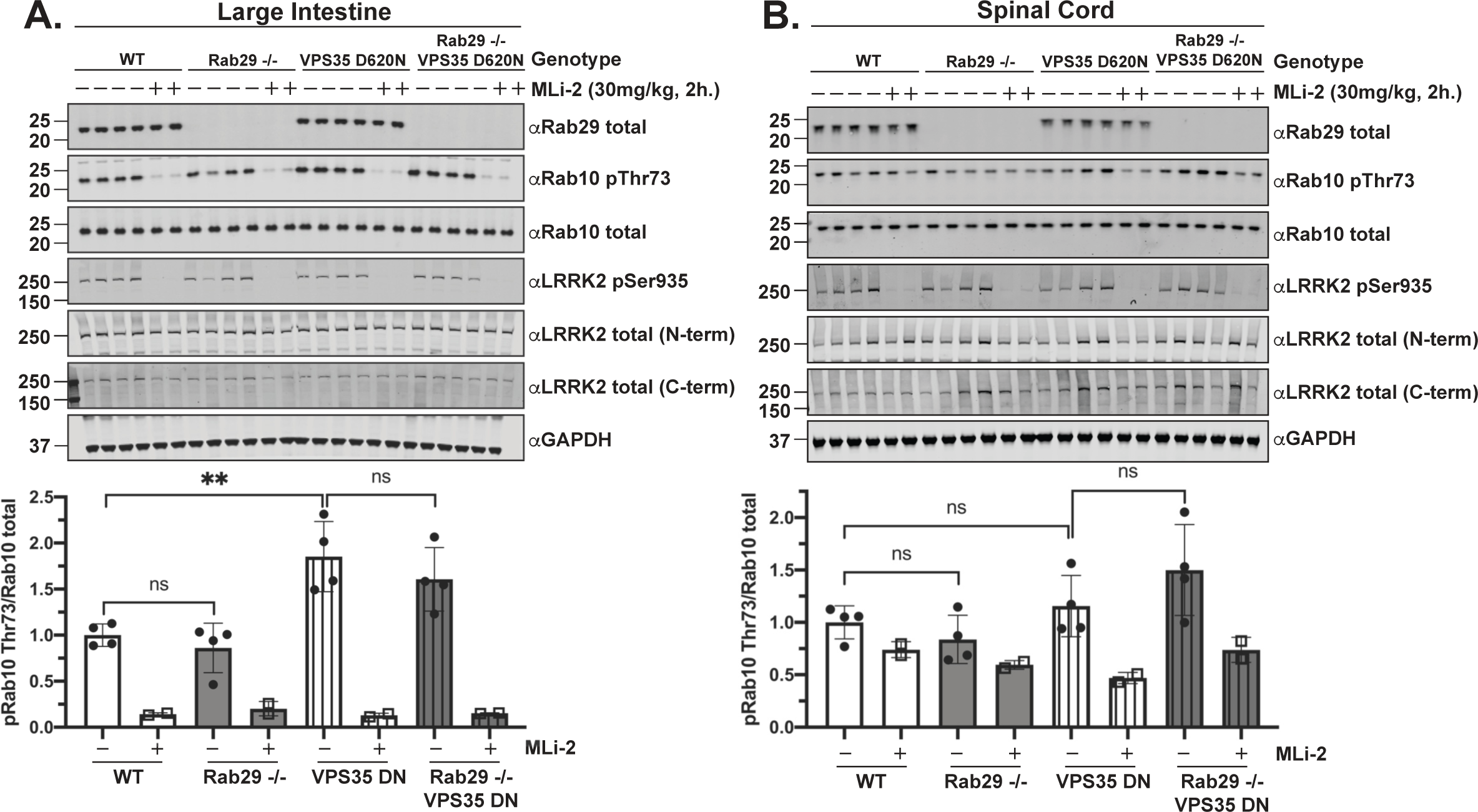
Rab29 knock-out does not reduce the enhanced LRRK2-mediated phosphorylation of Rab10 in VPS35[D620N] knock-in mice. **(A-B)** As in Figure 6 (B-D), the indicated 6-month-old matched mice were administered with vehicle (40% (w/v) (2-hydroxypropyl)-β-cyclodextrin) or 30 mg/kg MLi-2 dissolved in vehicle by subcutaneous injection 2 h prior to tissue collection. 40 μg of tissue extracts were subjected to quantitative immunoblot analysis with the indicated antibodies. The membranes were developed using the LI-COR Odyssey CLx Western Blot imaging system. Quantified data are presented as the ratios of phospho-Rab10/total Rab10, calculated using Image Studio software. Quantifications are presented as mean ± SD, normalized to vehicle treated, wildtype animals. Each lane represents a tissue sample from a different animal. Data were analyzed by one-way ANOVA with Tukey’s multiple comparisons test and there was a statistically significant difference between wildtype and VPS35[D620N] large intestine samples (**P = 0.0072 (A)), but not between wildtype and VPS35[D620N] spinal cord (P = 0.8768 (B)). All other comparisons were not statistically significant. Wildtype vs Rab29 knock-out: P = 0.9092 (A), P = 0.8629 (B). VPS35[D620N] vs Rab29 knock-out VPS35[D620N]: P = 0.6516 (A), 0.3935 (B).

**Supplementary Figure 7.**
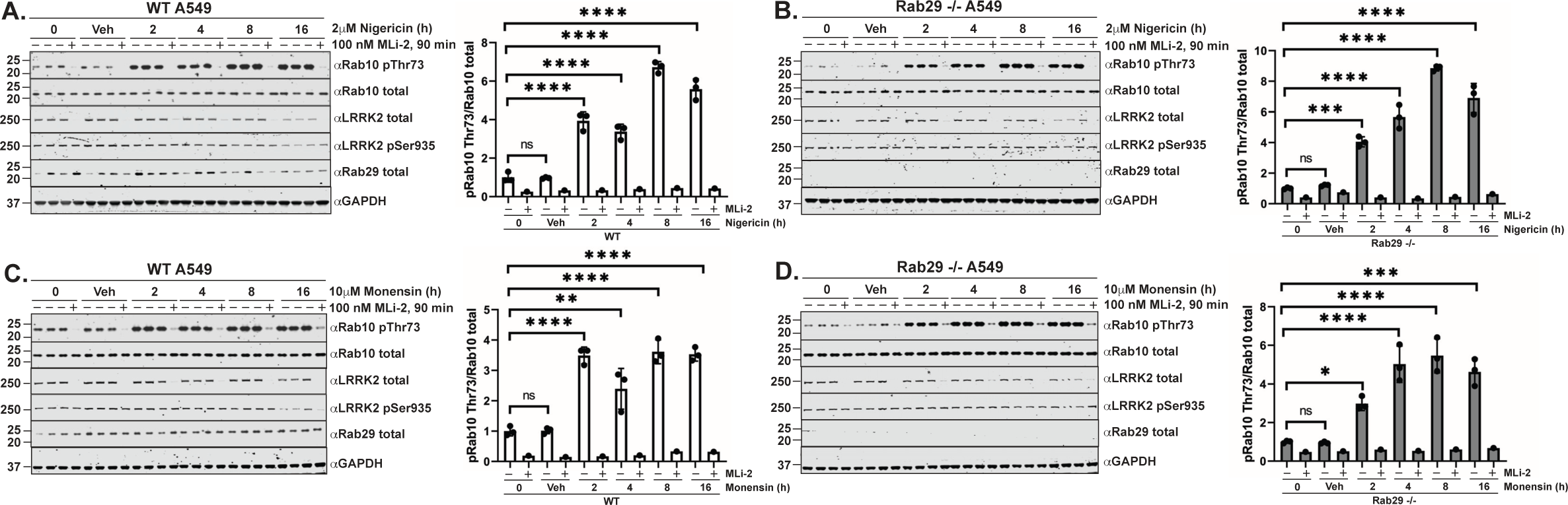
Cation ionophores nigericin and monensin enhance Rab10 phosphorylation in A549 wildtype and Rab29 knock-out cells. **(A-D)** A549 wildtype or Rab29 knock-out cells were treated with the appropriate vehicle or **(A-B)** 2 μM nigericin and **(C-D)** 10 μM monensin for the indicated periods of time. Cells were treated with DMSO or 100 nM LRRK2 inhibitor MLi-2 for 90 minutes prior to harvest. 15-20 μg of cell extract was subjected to quantitative immunoblot analysis with the indicated antibodies. The membranes were developed using the LI-COR Odyssey CLx Western Blot imaging system. Quantified data are presented as the ratios of phospho-Rab10/total Rab10, calculated using Image Studio software. Quantifications are presented as mean ± SD, normalized to the average of vehicle treated cells. Data were analyzed by one-way ANOVA with Dunnett’s multiple comparisons test and there was a statistically significant difference in LRRK2-mediated Rab10 phosphorylation in **(A)** between control and 2h treatment (****P < 0.0001), control and 4h treatment (****P < 0.0001), control and 8h treatment (****P < 0.0001), and control and 16h treatment (****P < 0.0001). There was a statistically significant difference in Rab10 phosphorylation in **(B)** between control and 2h treatment (***P = 0.0001), control and 4h treatment (****P < 0.0001), control and 8h treatment (****P < 0.0001), and control and 16h treatment (****P < 0.0001). There was a statistically significant difference in Rab10 phosphorylation in **(C)** between control and 2h treatment (****P < 0.0001), control and 4h treatment (**P = 0.0039), control and 8h treatment (****P < 0.0001), and control and 16h treatment (****P < 0.0001). There was a statistically significant difference in Rab10 phosphorylation in **(D)** between control and 2h treatment (*P = 0.0165), control and 4h treatment (****P < 0.0001), control and 8h treatment (****P < 0.0001), and control and 16h treatment (***P = 0.0001). Similar results were obtained in three independent experiments for each agonist.

## References

1 Zimprich, A., Biskup, S., Leitner, P., Lichtner, P., Farrer, M., Lincoln, S., Kachergus, J., Hulihan, M., Uitti, R. J., Calne, D. B., et al. (2004) Mutations in LRRK2 cause autosomal-dominant parkinsonism with pleomorphic pathology. Neuron 44, 601–607.

2 Paisán-Ruíz, C., Jain, S., Evans, E. W., Gilks, W. P., Simón, J., van der Brug, M., De Munain, A. L., Aparicio, S., Gil, A. M., Khan, N., et al. (2004) Cloning of the gene containing mutations that cause PARK8-linked Parkinson’s disease. Neuron 44, 595–600.

3 Taylor, M. and Alessi, D. R. (2020) Advances in elucidating the function of leucine-rich repeat protein kinase-2 in normal cells and Parkinson’s disease. Curr. Opin. Cell Biol., Elsevier Ltd 63, 102–113.

4 Domingo, A. and Klein, C. (2018) Genetics of Parkinson disease. Handb. Clin. Neurol. 1st ed., Elsevier B.V.

5 Price, A., Manzoni, C., Cookson, M. R. and Lewis, P. A. (2018) The LRRK2 signalling system. Cell Tissue Res., Springer Verlag 373, 39–50.

6 Jaleel, M., Nichols, R. J., Deak, M., Campbell, D. G., Gillardon, F., Knebel, A. and Alessi, D. R. (2007) LRRK2 phosphorylates moesin at threonine-558: Characterization of how Parkinson’s disease mutants affect kinase activity. Biochem. J. 405, 307–317.

7 Steger, M., Tonelli, F., Ito, G., Davies, P., Trost, M., Vetter, M., Wachter, S., Lorentzen, E., Duddy, G., Wilson, S., et al. (2016) Phosphoproteomics reveals that Parkinson’s disease kinase LRRK2 regulates a subset of Rab GTPases. Elife 5, e12813.

8 West, A. B., Moore, D. J., Biskup, S., Bugayenko, A., Smith, W. W., Ross, C. A., Dawson, V. L. and Dawson, T. M. (2005) Parkinson’s disease-associated mutations in leucine-rich repeat kinase 2 augment kinase activity. Proc. Natl. Acad. Sci. U. S. A. 102, 16842–16847.

9 Ito, G., Katsemonova, K., Tonelli, F., Lis, P., Baptista, M. A. S., Shpiro, N., Duddy, G., Wilson, S., Ho, P. W. L., Ho, S. L., et al. (2016) Phos-Tag analysis of Rab10 phosphorylation by LRRK2: A powerful assay for assessing kinase function and inhibitors. Biochem. J., Portland Press Ltd 473, 2671–2685.

10 Sheng, Z., Zhang, S., Bustos, D., Kleinheinz, T., Le Pichon, C. E., Dominguez, S. L., Solanoy, H. O., Drummond, J., Zhang, X., Ding, X., et al. (2012) Ser1292 autophosphorylation is an indicator of LRRK2 kinase activity and contributes to the cellular effects of PD mutations. Sci. Transl. Med. 4, 164ra161.

11 Lewis, P. A., Greggio, E., Beilina, A., Jain, S., Baker, A. and Cookson, M. R. (2007) The R1441C mutation of LRRK2 disrupts GTP hydrolysis. Biochem. Biophys. Res. Commun. 357, 668–671.

12 Li, X., Tan, Y. C., Poulose, S., Olanow, C. W., Huang, X. Y. and Yue, Z. (2007) Leucine-rich repeat kinase 2 (LRRK2)/PARK8 possesses GTPase activity that is altered in familial Parkinson’s disease R1441C/G mutants. J. Neurochem. 103, 238–247.

13 Liao, J., Wu, C. X., Burlak, C., Zhang, S., Sahm, H., Wang, M., Zhang, Z. Y., Vogel, K. W., Federici, M., Riddle, S. M., et al. (2014) Parkinson disease-associated mutation R1441H in LRRK2 prolongs the “active state” of its GTPase domain. Proc. Natl. Acad. Sci. U. S. A., National Academy of Sciences 111, 4055–4060.

14 Purlyte, E., Dhekne, H. S., Sarhan, A. R., Gomez, R., Lis, P., Wightman, M., Martinez, T. N., Tonelli, F., Pfeffer, S. R. and Alessi, D. R. (2018) Rab29 activation of the Parkinson’s disease-associated LRRK2 kinase. EMBO J 37, 1–18.

15 Beilina, A., Rudenko, I. N., Kaganovich, A., Civiero, L., Chau, H., Kalia, S. K., Kalia, L. V, Lobbestael, E., Chia, R., Ndukwe, K., et al. (2014) Unbiased screen for interactors of leucinerich repeat kinase 2 supports a common pathway for sporadic and familial Parkinson disease. Proc Natl Acad Sci U S A 111, 2626–2631.

16 Liu, Z., Bryant, N., Kumaran, R., Beilina, A., Abeliovich, A., Cookson, M. R. and West, A. B. (2018) LRRK2 phosphorylates membrane-bound Rabs and is activated by GTP-bound Rab7L1 to promote recruitment to the trans-Golgi network. Hum Mol Genet 27, 385–395.

17 Fujimoto, T., Kuwahara, T., Eguchi, T., Sakurai, M., Komori, T. and Iwatsubo, T. (2018) Parkinson’s disease-associated mutant LRRK2 phosphorylates Rab7L1 and modifies trans-Golgi morphology. Biochem Biophys Res Commun 495, 1708–1715.

18 Watanabe, R., Buschauer, R., Böhning, J., Audagnotto, M., Lasker, K., Wen Lu, T., Boassa, D., Taylor, S. S. and Villa, E. (2020) The In situ Structure of Parkinson’s Disease-Linked LRRK2. Biophys. J. 118, 1508–1518.

19 Deniston, C. K., Salogiannis, J., Mathea, S., Snead, D. M., Lahiri, I., Matyszewski, M., Donosa, O., Watanabe, R., Böhning, J., Shiau, A. K., et al. (2020) Structure of LRRK2 in Parkinson’s disease and model for microtubule interaction. Nature.

20 Schmidt, S. H., Knape, M. J., Boassa, D., Mumdey, N., Kornev, A. P., Ellisman, M. H., Taylor, S. S. and Herberg, F. W. (2019) The dynamic switch mechanism that leads to activation of LRRK2 is embedded in the DFGψ motif in the kinase domain. Proc. Natl. Acad. Sci. U. S. A., National Academy of Sciences 116, 14979–14988.

21 Mir, R., Tonelli, F., Lis, P., Macartney, T., Polinski, N. K., Martinez, T. N., Chou, M. Y., Howden, A. J. M., Konig, T., Hotzy, C., et al. (2018) The Parkinson’s disease VPS35[D620N] mutation enhances LRRK2-mediated Rab protein phosphorylation in mouse and human. Biochem J 475, 1861–1883.

22 Doggett, E. A., Zhao, J., Mork, C. N., Hu, D. and Nichols, R. J. (2012) Phosphorylation of LRRK2 serines 955 and 973 is disrupted by Parkinson’s disease mutations and LRRK2 pharmacological inhibition. J. Neurochem. 120, 37–45.

23 Dzamko, N., Deak, M., Hentati, F., Reith, A. D., Prescott, A. R., Alessi, D. R. and Nichols, R. J. (2010) Inhibition of LRRK2 kinase activity leads to dephosphorylation of Ser 910/Ser935, disruption of 14-3-3 binding and altered cytoplasmic localization. Biochem. J. 430, 405–413.

24 Alessi, D. R. and Sammler, E. (2018) LRRK2 kinase in Parkinson’s disease. Science (80-.)., American Association for the Advancement of Science 360, 36–37.

25 Tolosa, E., Vila, M., Klein, C. and Rascol, O. (2020) LRRK2 in Parkinson disease: challenges of clinical trials. Nat. Rev. Neurol., Springer US 16, 97–107.

26 Steger, M., Diez, F., Dhekne, H. S., Lis, P., Nirujogi, R. S., Karayel, O., Tonelli, F., Martinez, T. N., Lorentzen, E., Pfeffer, S. R., et al. (2017) Systematic proteomic analysis of LRRK2-mediated Rab GTPase phosphorylation establishes a connection to ciliogenesis. Elife 6, e31012.

27 Pfeffer, S. R. (2017) Rab GTPases: Master regulators that establish the secretory and endocytic pathways. Mol. Biol. Cell, American Society for Cell Biology 28, 712–715.

28 Jeong, G. R., Jang, E. H., Bae, J. R., Jun, S., Kang, H. C., Park, C. H., Shin, J. H., Yamamoto, Y., Tanaka-Yamamoto, K., Dawson, V. L., et al. (2018) Dysregulated phosphorylation of Rab GTPases by LRRK2 induces neurodegeneration. Mol. Neurodegener., BioMed Central Ltd. 13, 8.

29 Gomez, R. C., Wawro, P., Lis, P., Alessi, D. R. and Pfeffer, S. R. (2019) Membrane association but not identity is required for LRRK2 activation and phosphorylation of Rab GTPases. J. Cell Biol., NLM (Medline) 218, 4157–4170.

30 Waschbüsch, D., Purlyte, E., Pal, P., McGrath, E., Alessi, D. R. and Khan, A. R. (2020) Structural Basis for Rab8a Recruitment of RILPL2 via LRRK2 Phosphorylation of Switch 2. Structure, Cell Press 28, 406–417.

31 Dhekne, H. S., Yanatori, I., Gomez, R. C., Tonelli, F., Diez, F., Schüle, B., Steger, M., Alessi, D. R. and Pfeffer, S. R. (2018) A pathway for parkinson’s disease LRRK2 kinase to block primary cilia and sonic hedgehog signaling in the brain. Elife, eLife Sciences Publications Ltd 7.

32 Sobu, Y., Wawro, P. S., Dhekne, H. S. and Pfeffer, S. R. (2020) Pathogenic LRRK2 regulates ciliation probability upstream of Tau Tubulin kinase 2. bioRxiv, Cold Spring Harbor Laboratory 2020.04.07.029983.

33 Berndsen, K., Lis, P., Yeshaw, W. M., Wawro, P. S., Nirujogi, R. S., Wightman, M., Macartney, T., Dorward, M., Knebel, A., Tonelli, F., et al. (2019) PPM1H phosphatase counteracts LRRK2 signaling by selectively dephosphorylating rab proteins. Elife, eLife Sciences Publications Ltd 8.

34 Lill, C. M., Roehr, J. T., McQueen, M. B., Kavvoura, F. K., Bagade, S., Schjeide, B. M., Schjeide, L. M., Meissner, E., Zauft, U., Allen, N. C., et al. (2012) Comprehensive research synopsis and systematic meta-analyses in Parkinson’s disease genetics: The PDGene database. PLoS Genet 8, e1002548.

35 Tucci, A., Nalls, M. A., Houlden, H., Revesz, T., Singleton, A. B., Wood, N. W., Hardy, J. and Paisan-Ruiz, C. (2010) Genetic variability at the PARK16 locus. Eur J Hum Genet 18, 1356–1359.

36 Simón-Sánchez, J., Schulte, C., Bras, J. M., Sharma, M., Gibbs, J. R., Berg, D., Paisan-Ruiz, C., Lichtner, P., Scholz, S. W., Hernandez, D. G., et al. (2009) Genome-wide association study reveals genetic risk underlying Parkinson’s disease. Nat. Genet., Nature Publishing Group 41, 1308–1312.

37 Satake, W., Nakabayashi, Y., Mizuta, I., Hirota, Y., Ito, C., Kubo, M., Kawaguchi, T., Tsunoda, T., Watanabe, M., Takeda, A., et al. (2009) Genome-wide association study identifies common variants at four loci as genetic risk factors for Parkinson’s disease. Nat. Genet., Nature Publishing Group 41, 1303–1307.

38 Nalls, M. A., Pankratz, N., Lill, C. M., Do, C. B., Hernandez, D. G., Saad, M., DeStefano, A. L., Kara, E., Bras, J., Sharma, M., et al. (2014) Large-scale meta-analysis of genome-wide association data identifies six new risk loci for Parkinson’s disease. Nat Genet 46, 989–993.

39 MacLeod, D. A., Rhinn, H., Kuwahara, T., Zolin, A., Di Paolo, G., MacCabe, B. D., Marder, K. S., Honig, L. S., Clark, L. N., Small, S. A., et al. (2013) RAB7L1 Interacts with LRRK2 to Modify Intraneuronal Protein Sorting and Parkinson’s Disease Risk. Neuron, Elsevier 77, 425–439.

40 Pihlstrøm, L., Rengmark, A., Bjørnarå, K. A., Dizdar, N., Fardell, C., Forsgren, L., Holmberg, B., Larsen, J. P., Linder, J., Nissbrandt, H., et al. (2015) Fine mapping and resequencing of the PARK16 locus in Parkinson’s disease. J. Hum. Genet. 60, 357–362.

41 McGrath, E., Waschbüsch, D., Baker, B. M. and Khan, A. R. (2019) LRRK2 binds to the Rab32 subfamily in a GTP-dependent manner via its armadillo domain. Small GTPases, Taylor & Francis 1–14.

42 Dodson, M. W., Zhang, T., Jiang, C., Chen, S. and Guo, M. (2012) Roles of the Drosophila LRRK2 homolog in Rab7-dependent lysosomal positioning. Hum. Mol. Genet. 21, 1350–1363.

43 Kuwahara, T., Inoue, K., D’Agati, V. D., Fujimoto, T., Eguchi, T., Saha, S., Wolozin, B., Iwatsubo, T. and Abeliovich, A. (2016) LRRK2 and RAB7L1 coordinately regulate axonal morphology and lysosome integrity in diverse cellular contexts. Sci Rep 6, 29945.

44 Mazza, M. C., Nguyen, V., Beilina, A., Ding, J. and Cookson, M. R. (2020) Combined knockout of Lrrk2 and Rab29 does not result in behavioral abnormalities in vivo. bioRxiv, Cold Spring Harbor Laboratory 2020.05.13.093708.

45 Wang, S., Ma, Z., Xu, X., Wang, Z., Sun, L., Zhou, Y., Lin, X., Hong, W. and Wang, T. (2014) A role of Rab29 in the integrity of the trans-Golgi network and retrograde trafficking of mannose-6-phosphate receptor. PLoS One 9, e96242.

46 Beilina, A., Bonet-Ponce, L., Kumaran, R., Kordich, J. J., Ishida, M., Mamais, A., Kaganovich, A., Saez-Atienzar, S., Gershlick, D. C., Roosen, D. A., et al. (2020) The Parkinson’s Disease Protein LRRK2 Interacts with the GARP Complex to Promote Retrograde Transport to the trans-Golgi Network. Cell Rep. 31, 107614.

47 Spanò, S., Liu, X. and Galán, J. E. (2011) Proteolytic targeting of Rab29 by an effector protein distinguishes the intracellular compartments of human-adapted and broad-host Salmonella. Proc. Natl. Acad. Sci. U. S. A., National Academy of Sciences 108, 18418–18423.

48 Wasmeier, C., Romao, M., Plowright, L., Bennett, D. C., Raposo, G. and Seabra, M. C. (2006) Rab38 and Rab32 control post-Golgi trafficking of melanogenic enzymes. J Cell Biol 175, 271–281.

49 Fell, M. J., Mirescu, C., Basu, K., Cheewatrakoolpong, B., DeMong, D. E., Ellis, J. M., Hyde, L. A., Lin, Y., Markgraf, C. G., Mei, H., et al. (2015) MLi-2, a potent, selective, and centrally active compound for exploring the therapeutic potential and safety of LRRK2 kinase inhibition. J. Pharmacol. Exp. Ther. 355, 397–409.

50 Spieker-Polet, H., Sethupathi, P., Yam, P. C. and Knight, K. L. (1995) Rabbit monoclonal antibodies: Generating a fusion partner to produce rabbit-rabbit hybridomas. Proc. Natl. Acad. Sci. U. S. A. 92, 9348–9352.

51 Lis, P., Burel, S., Steger, M., Mann, M., Brown, F., Diez, F., Tonelli, F., Holton, J. L., Ho, P. W., Ho, S. L., et al. (2018) Development of phospho-specific Rab protein antibodies to monitor in vivo activity of the LRRK2 Parkinson’s disease kinase. Biochem. J. 475, 1–22.

52 Liang, Y., Lin, S., Zou, L., Zhou, H., Zhang, J., Su, B. and Wan, Y. (2012) Expression profiling of Rab GTPases reveals the involvement of Rab20 and Rab32 in acute brain inflammation in mice. Neurosci Lett 527, 110–114.

53 Livak, K. J. and Schmittgen, T. D. (2001) Analysis of relative gene expression data using real-time quantitative PCR and the 2-ΔΔCT method. Methods 25, 402–408.

54 Kluss, J. H., Conti, M. M., Kaganovich, A., Beilina, A., Melrose, H. L., Cookson, M. R. and Mamais, A. (2018) Detection of endogenous S1292 LRRK2 autophosphorylation in mouse tissue as a readout for kinase activity. npj Park. Dis., Springer US 4, 1–5.

55 Wiggin, G. R., Soloaga, A., Foster, J. M., Murray-Tait, V., Cohen, P. and Arthur, J. S. C. (2002) MSK1 and MSK2 Are Required for the Mitogen-and Stress-Induced Phosphorylation of CREB and ATF1 in Fibroblasts. Mol. Cell. Biol. 22, 2871–2881.

56 Tian, X., Azpurua, J., Hine, C., Vaidya, A., Myakishev-Rempel, M., Ablaeva, J., Mao, Z., Nevo, E., Gorbunova, V. and Seluanov, A. (2013) High-molecular-mass hyaluronan mediates the cancer resistance of the naked mole rat. Nature 499, 346–349.

57 Eguchi, T., Kuwahara, T., Sakurai, M., Komori, T., Fujimoto, T., Ito, G., Yoshimura, S. I., Harada, A., Fukuda, M., Koike, M., et al. (2018) LRRK2 and its substrate Rab GTPases are sequentially targeted onto stressed lysosomes and maintain their homeostasis. Proc Natl Acad Sci U S A 115, E9115–E9124.

58 Fan, Y., Howden, A. J. M., Sarhan, A. R., Lis, P., Ito, G., Martinez, T. N., Brockmann, K., Gasser, T., Alessi, D. R. and Sammler, E. M. (2018) Interrogating Parkinson’s disease LRRK2 kinase pathway activity by assessing Rab10 phosphorylation in human neutrophils. Biochem J 475, 23–44.

59 Kelly, K., Wang, S., Boddu, R., Liu, Z., Moukha-Chafiq, O., Augelli-Szafran, C. and West, A. B. (2018) The G2019S mutation in LRRK2 imparts resiliency to kinase inhibition. Exp. Neurol., Academic Press Inc. 309, 1–13.

60 Casola, S. (2010) Mouse models for miRNA expression: the ROSA26 locus. Methods Mol. Biol. (Methods Protoc., Humana Press, Totowa, NJ 667, 145–163.

61 Nichols, R. J., Dzamko, N., Morrice, N. A., Campbell, D. G., Deak, M., Ordureau, A., Macartney, T., Tong, Y., Shen, J., Prescott, A. R., et al. (2010) 14-3-3 Binding to LRRK2 is disrupted by multiple Parkinson’s disease-associated mutations and regulates cytoplasmic localization. Biochem. J. 430, 393–404.

62 Steinrauf, L. K. and Pinkerton, M. (1968) The Structure of Nigericin. Biochem Biophys Res Commun 33, 29–31.

63 Tartakoff, A. M. (1983) Perturbation of vesicular traffic with the carboxylic ionophore monensin. Cell 32, 1026–1028.

64 Mauthe, M., Orhon, I., Rocchi, C., Zhou, X., Luhr, M., Hijlkema, K. J., Coppes, R. P., Engedal, N., Mari, M. and Reggiori, F. (2018) Chloroquine inhibits autophagic flux by decreasing autophagosome-lysosome fusion. Autophagy, Taylor & Francis 14, 1435–1455.

65 Cirman, T., Oreŝic, K., Mazovec, G. D., Turk, V., Reed, J. C., Myers, R. M., Salvesen, G. S. and Turk, B. (2004) Selective Disruption of Lysosomes in HeLa Cells Triggers Apoptosis Mediated by Cleavage of Bid by Multiple Papain-like Lysosomal Cathepsins. J. Biol. Chem. 279, 3578–3587.

66 Herbst, S., Campbell, P., Harvey, J., Bernard, E. M., Papayannopoulos, V., Wood, N. W., Morris, H. R. and Gutierrez, M. G. (2020) LRRK2 activation controls the repair of damaged endomembranes in macrophages. EMBO J. 39, e104494.

67 Kuwahara, T., Funakawa, K., Komori, T., Sakurai, M., Yoshii, G., Eguchi, T., Fukuda, M. and Iwatsubo, T. (2020) Roles of lysosomotropic agents on LRRK2 activation and Rab10 phosphorylation. Neurobiol. Dis., Elsevier 145, 105081.

68 Bonet-Ponce, L., Beilina, A., Williamson, C. D., Lindberg, E., Kluss, J. H., Saez-Atienzar, S., Landeck, N., Kumaran, R., Mamais, A., Bleck, C. K. E., et al. (2020) LRRK2 mediates tubulation and vesicle sorting from membrane damaged lysosomes. bioRxiv 2020.01.23.917252.

69 Waschbüsch, D., Michels, H., Strassheim, S., Ossendorf, E., Kessler, D., Gloeckner, C. J. and Barnekow, A. (2014) LRRK2 transport is regulated by its novel interacting partner Rab32. PLoS One 9, e111632.

70 Kluss, J. H., Mazza, M. C., Li, Y., Manzoni, C., Lewis, P. A., Cookson, M. R. and Mamais, A. (2020) Preclinical Modeling of Chronic Inhibition of the Parkinson’s Disease Associated Kinase LRRK2 Reveals Altered Function of the Endolysosomal System in Vivo. Mol. Neurodegener. 1–27.

71 Biskup, S., Moore, D. J., Celsi, F., Higashi, S., West, A. B., Andrabi, S. A., Kurkinen, K., Yu, S. W., Savitt, J. M., Waldvogel, H. J., et al. (2006) Localization of LRRK2 to membranous and vesicular structures in mammalian brain. Ann. Neurol. 60, 557–569.

72 Kett, L. R., Boassa, D., Ho, C. C. Y., Rideout, H. J., Hu, J., Terada, M., Ellisman, M. and Dauer, W. T. (2012) LRRK2 Parkinson disease mutations enhance its microtubule association. Hum. Mol. Genet. 21, 890–899.

73 Madero-Pérez, J., Fernández, B., Lara Ordóñez, A. J., Fdez, E., Lobbestael, E., Baekelandt, V. and Hilfiker, S. (2018) RAB7L1-mediated relocalization of LRRK2 to the golgi complex causes centrosomal deficits via RAB8A. Front. Mol. Neurosci. 11, 1–19.

74 Bae, E. J., Kim, D. K., Kim, C., Mante, M., Adame, A., Rockenstein, E., Ulusoy, A., Klinkenberg, M., Jeong, G. R., Bae, J. R., et al. (2018) LRRK2 kinase regulates α-synuclein propagation via RAB35 phosphorylation. Nat. Commun., Springer US 9.

